# A multidimensional atlas of human glioblastoma organoids reveals highly coordinated molecular networks and effective drugs

**DOI:** 10.1101/2023.01.24.525374

**Authors:** Changwen Wang, Meng Sun, Chunxuan Shao, Lisa Schlicker, Yue Zhuo, Yassin Harim, Tianping Peng, Weili Tian, Nadja Stöffler, Martin Schneider, Dominic Helm, Jan-Philipp Mallm, Yonghe Wu, Almut Schulze, Hai-Kun Liu

## Abstract

Recent advances in the genomics of glioblastoma (GBM) led to the introduction of molecular neuropathology but failed to translate into treatment improvement. This is largely attributed to the genetic and phenotypic heterogeneity of GBM, which are considered the major obstacle to GBM therapy. Here, we use advanced human GBM organoid (LEGO: <u>L</u>aboratory <u>E</u>ngineered <u>G</u>lioblastoma <u>O</u>rganoid) and provide an unprecedented comprehensive characterization of LEGO models using single-cell transcriptome, DNA methylome, metabolome, lipidome, proteome, and phospho-proteome analysis. We discovered that genetic heterogeneity dictates functional heterogeneity across molecular layers and demonstrates that *NF1* mutation drives mesenchymal signature. Most importantly, we found that glycerol lipid reprogramming is a hallmark of GBM, and several targets and drugs were discovered along this line. We also provide a genotype-based drug reference map using LEGO-based drug screen. This study provides novel human GBM models and a research path toward effective GBM therapy.

## Introduction

Oncogenic genetic alteration is a fundamental hallmark of human cancers and has been utilized to characterize genotype-specific molecular features, which form the basis for personalized treatment of cancer patients ^1,2^. Based on these efforts, genotype-based personalized cancer treatment options are already available for many human cancers, i.e., breast cancer, lung cancer, and leukemia ^1^. However, it remains challenging to expand personalized treatment to most cancer patients ^3^.

GBM is the most malignant type of primary brain cancer and was one of the first tumor entities selected for The Cancer Genome Atlas (TCGA) project ^4,5^. With the continuous efforts in genomic analysis of GBM, it has been suggested that GBM is a heterogeneous group of diseases of different molecular subtypes based on RNA expression, DNA methylation, or recently via multi-omics analysis ^4,6–8^. Single-cell RNA-sequencing (scRNA-seq) analysis of human GBM identified intratumoral heterogeneity of GBM, which provides a single-cell molecular description of human GBM, it was suggested that the GBM cells are of high plasticity, which may switch among the molecular phenotypes ^9,10^. However, it must be noted that how tumor genotype contributes to the molecular phenotype-related plasticity remains unclear, i.e., *NF1* mutation in human GBM is associated with a mesenchymal feature, but this has not been verified in animal models ^4,5^. And it is much more challenging to perform in-depth single-cell DNA sequencing. In contrast to the rapid development of molecular characterization of GBM, the clinical treatment options for human GBM patients remain to be neurosurgery, plus radiotherapy and temozolomide(TMZ)-based chemotherapy ^11^. There is a clear gap between the comprehensive molecular description of GBM and treatment improvement, which need to be highly prioritized for future GBM research.

A genome-based personalized treatment of cancer patients requires a solid understanding of genotype-specific cancer pathway dependency and actionable target identification. Model systems of GBM have been utilized to systematically analyze and compare differences in cancer cells with different mutation combinations. Genetically modified mouse models have been used to determine the function of selected genes and identify the cell of origin in brain tumors ^12^. However, mouse models often do not represent the molecular pathology of human tumors ^13^. Patient-derived xenograft (PDX) models or organoids harbor patient tumor cells. Still, they are limited by complex genetic background variations, differences in treatment histories, and, most importantly, the lack of suitable controls ^14^. Most importantly, the tumor growth characteristics identified in PDX models were found to be more dependent on the mouse strain than tumor type ^15^, suggesting the PDX model may generate many artificial readouts irrelevant to primary human tumors.

The recent development of organoid technology coupled with gene editing by CRISPR/Cas9 allows the rapid generation of genetic mutations in human-derived tissues to model cancer progression ^16^. Initial attempts were made using induced pluripotent stem cells (iPSCs)-derived cerebral organoids to generate glioma-like organoids ^17,18^. This model provides the opportunity to develop genetically customized GBM models derived from single iPSC clones. Therefore, a rigorous follow-up analysis can be performed using this experimental system.

Here we generated a set of iPSC-based human GBM organoid models (**LEGO**: Laboratory Engineered Glioblastoma Organoid) based on CRISPR/Cas9 engineered loss of tumor suppressors, which are frequently mutated in human GBM patients. Comprehensive analysis of LEGOs demonstrates their great potential in identifying novel molecular features in cancer cells, providing a path toward personalized treatment of human GBM.

## Results

### Generation of LEGOs with defined genetic mutations

We used human iPSC-derived organoids to dissect the functional consequences of genetic heterogeneity in GBM (Figure 1A). Using CRISPR/Cas9, we generated a spectrum of mutation combinations (**PT**: *<u>P</u>TEN^-/-^*; *<u>T</u>P53^-/-^*, **PTCC**: *<u>P</u>TEN^-/-^*; *<u>T</u>P53^-/-^*; *<u>C</u>DKN2A^-/-^*; *<u>C</u>DKN2B^-/-^*, **PTN**: *<u>P</u>TEN^-/-^*; *<u>T</u>P53^-/-^*; *<u>N</u>F1^-/-^*), which are among the most frequently mutated tumor suppressors in GBM patients ^5^, in wildtype (WT) iPSCs express GFP. The knockout of individual genes was confirmed by Western blotting and sequencing (Figures S1A, S1B). All iPSCs clones grew well except that the PTN clone showed signs of differentiation, which was reported previously and could be controlled by MEK inhibitor PD0325901 ^19^. These iPSCs were then differentiated into organoids with a previously described cerebral organoid protocol ^20^. Although starting from the same number of cells, all the LEGOs grew faster and more extensively than WT organoids (Figure 1B, 1C), indicating the activation of cell proliferation and growth pathways following the oncogenic mutations. Interestingly, the size of PT organoids was the biggest among the three mutant groups (Figure 1C).

**Figure 1.**
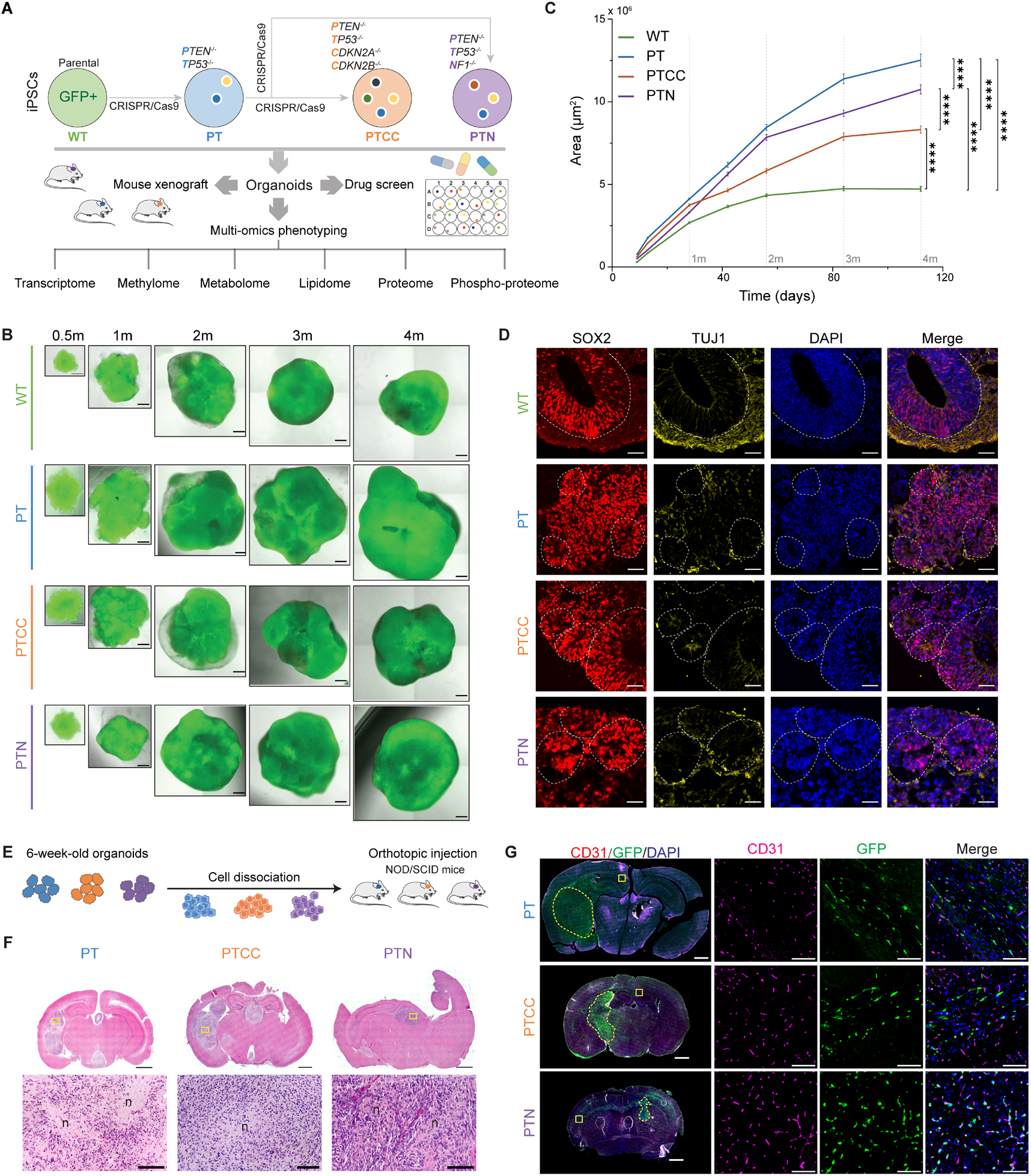
Generation and histological characterization of LEGOs with defined genetic mutations. (A). Schematic illustration of experimental procedures in this study. (B). Representative images showing the morphology of organoids at different ages. (C). Organoid growth curves. *P* values of the comparison between different groups of organoids were calculated by Two-way ANOVA. Data are represented as mean ± SEM for quantifying the 2D area of at least 65 organoids from at least four independent batches at each time point. (D). Representative immunofluorescent staining images of 1-month-old organoids stained with SOX2 and TUJ1. White circles show rosette-like structures. Scale bars, 50 μm. (E). Schematic diagram illustrating mouse xenograft workflow. (F). Representative H&E staining images of brain tumors in mouse xenografts. “n” in the enlarged image marks necrotic areas. Scale bar, 1000 μm for the overview, and 100 μm for insets (G). Representative immunofluorescent images of the tumor-infiltrating area stained with GFP and CD31. Note a strong association between GFP (green) and CD31(magenta) -positive cells in the PTN xenografts. Scale bars, 1000 μm for overview images, and 100 μm for insets. See also Figure S1

The histological analysis showed that the LEGOs exhibited similar structures compared to the WT organoids, indicated by the expression of SOX2 (SRY-box transcription factor 2) and TUJ1 (neuron-specific class III beta-tubulin) (Figure 1D). However, all mutant LEGOs show an increased stem/progenitor population (Figure 1D). H&E staining revealed nuclear atypia in LEGOs after more than one month of culture (Figure S1C), indicating signs of malignant transformation. To investigate whether the LEGOs are tumorigenic *in vivo,* we performed xenograft experiments, as illustrated in Figure 1E. All LEGOs initiated fatal brain tumors upon xenograft (Figure 1F, S1D). The grafted GFP^+^ LEGO cells showed infiltrative and angiogenic growth patterns (Figure 1G, S1E). Moreover, PTN xenografts exhibited a more infiltrative growth pattern with tumor cells migrating to the other hemisphere and being tightly associated with blood vessels (Figure 1G, S1E), suggesting that loss of *NF1* results in a more invasive phenotype, which is a feature of the mesenchymal molecular phenotype of human GBM. In addition, all grafts expressed markers, like astrocyte marker GFAP (glial fibrillary acid protein), neural stem cell marker Nestin, and cell proliferation marker Ki67 (Figure S1F), the tumors also show signs of necrosis, which is a hallmark for GBM (Figure 1F). These results demonstrate that the LEGO organoids are GBM organoids.

### ScRNA-seq analysis reveals shared and genotype-specific alterations during early tumor development

One of the advantages of cerebral organoids is that they contain heterogeneous neural cell populations and maintain differentiation hierarchies ^20^, thus can be used to study cellular heterogeneity and plasticity. To fully characterize the LEGOs on the single-cell level and to understand how different genetic mutations affect cellular heterogeneity, we performed scRNA-seq on one- and four-month-old LEGOs. In total, we obtained results from 70617 cells for further analysis.

We next performed UMAP (uniform manifold approximation and projection) analysis to visualize cell differentiation trajectory ^21^. UMAP of one-month-old LEGOs show two major lineages (neuron and astrocyte), which was confirmed by the expression of immature neuronal marker DCX (doublecortin), astrocytic marker FABP7 (fatty acid binding protein 7) and APOE (apolipoprotein E), and neural stem/progenitor marker SOX2 (Figure 2A, S2A). The one-month-old WT organoids mainly differentiated toward the neuronal lineage, whereas the PT and PTCC organoids switched to astrocytic differentiation (Figure 2A, S2A). The PTN organoids exhibited limited neuronal differentiation and reduced astrocytic differentiation (Figure 2A, S2A), suggesting a general blockage of neural differentiation. We also observed increased expression of neural stem/progenitor markers like SOX2 in all the LEGOs, indicating differentiation blockage upon loss of tumor suppressors, consistent with staining (Figure 1D, S2A). Interestingly, PTCC organoids highly express WNT regulators in the glial progenitor population, suggesting the activation of the WNT pathway upon loss of CDKN2A/2B (Figure 2B). Surprisingly, the PTN organoids activate several HOX transcription factors in the stem cell clusters (Figure 2C). The HOX genes have been reported to be involved in the induction of EMT (epithelial-mesenchymal transition) in other cancers ^22^, and they are not expressed in normal neural cells (Figure S2B), which highly suggests that PTN organoids may activate a non-neural transcriptional program to acquire a more aggressive phenotype.

**Figure 2.**
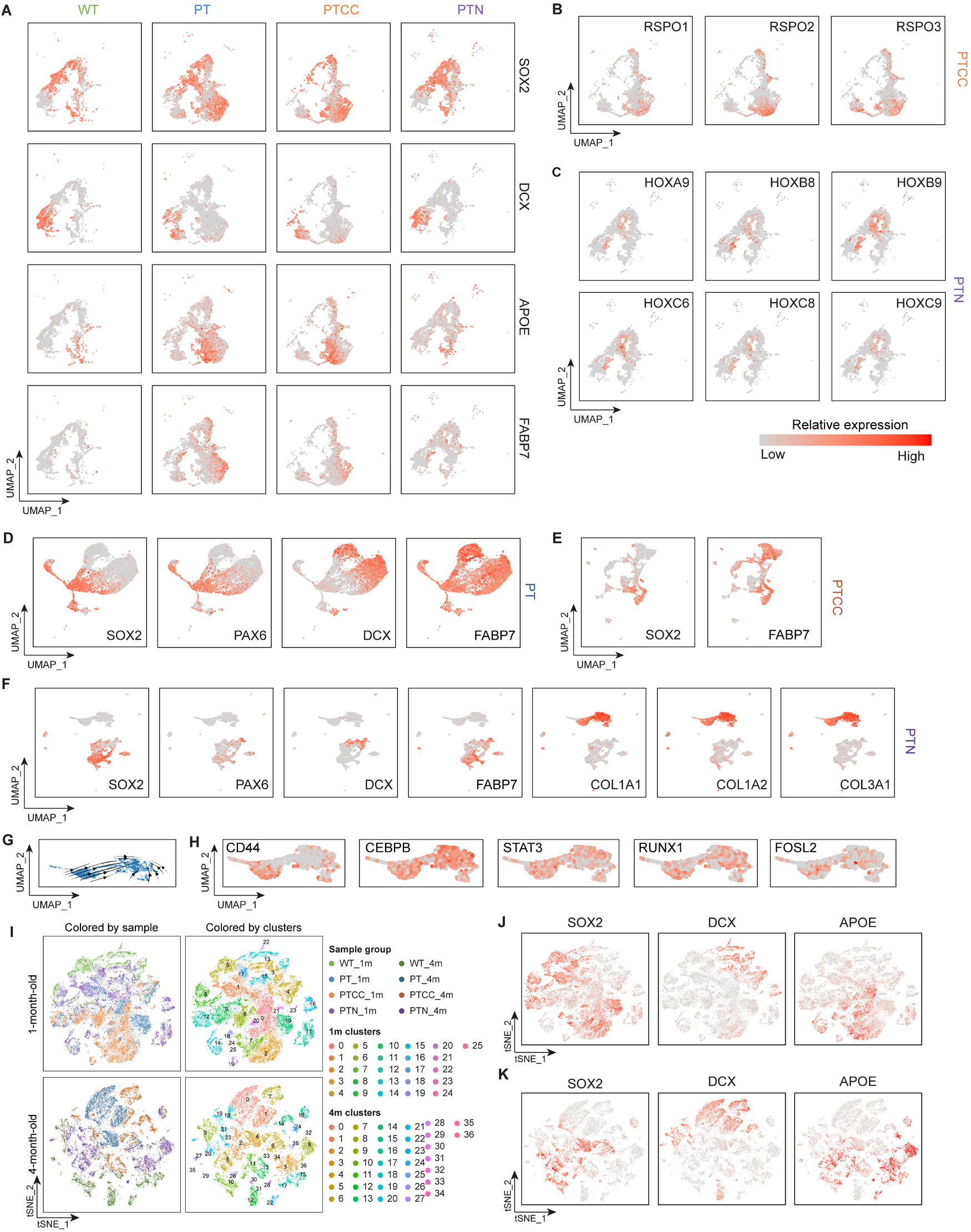
ScRNA-seq analysis reveals shared and genotype-specific alterations during early GBM development. (A). UMAP plots show the relative expression of lineage markers in one-month-old organoids. (B). UMAP plots for RSPO genes in one-month-old PTCC organoids. (C). UMAP plots for HOX genes in one-month-old PTN organoids. UMAP plots for lineage markers in four-month-old PT (D), PTCC (E), and PTN (F) organoids. (G). RNA velocity analysis of the mesenchymal-like clusters in PTN organoids. (H). UMAP plots of the mesenchymal-related marker genes in PTN organoids. (I). t-SNE plots of the one- and four-month-old organoids colored by sample groups or clusters. t-SNE plots for lineage markers in one- (J) and four-month-old (K) organoids. See also Figure S2.

The scRNA-seq results from four-month-old organoids demonstrate that the PT organoids are dominated by two major cell populations, one shows high expression of stem/progenitor cell markers like SOX2 and PAX6 (paired box 6), and the other express the immature neuron marker DCX and the astrocyte marker FABP7 (Figure 2D), indicating a proneural-like tumor cell feature. PTCC organoids also maintain a differentiation trajectory towards the FABP7 astrocytic lineage from the SOX2-positive stem cell cluster (Figure 2E). Strikingly, there are two major differentiation lineages in PTN organoids; one is the neural lineage, as indicated by the expression of SOX2, PAX6, and DCX (Figure 2F), while the other lineage highly expresses collagen genes and can be divided into two clusters (Figure 2F). The RNA velocity analysis suggested a possible differentiation hierarchy between the two clusters (Figure 2G), with the stem-cell-like cluster expressing CD44 (Figure 2H). Moreover, this lineage was positive for mesenchymal master regulators like STAT3 (signal transducer and activator of transcription 3), C/EBPB (CCAAT enhancer binding protein beta), RUNX1 (RUNX family transcription factor 1), and FOSL2 (FOS like 2, AP-1 transcription factor subunit) (Figure 2H) ^23^. We also found that these cells express unique markers like PAX7 (paired box 7) and CHODL (chondrolectin) (Figure S2C), which can potentially be used to identify these cells in human cancers.

Next, we annotated the cell clusters using reference gene signatures derived from GBM patient single-cell transcriptome data ^24,25^. The LEGOs contain major tumor cell populations such as “stem-like”, “proliferating stem-like”, and “differentiated-like” cells (Figure S2D). Moreover, PT was dominated by the “stem-like” cell population resembling the proneural subtype (Figure S2E) ^24^. PTN showed an increased proportion of “differentiated-like” cells mimicking the mesenchymal subtype (Figure S2E) ^24^. Moreover, we calculated the single cell meta score ^9^ of the mutant organoids and found that PT organoids were dominated by the NPC-like cells and PTN organoids were dominated by the MES-like cells (Figure S2F). These results suggest that *NF1* mutation drives a mesenchymal-like lineage during organoid development, and it will be interesting to trace the origin of these cells in the future.

The LEGO models recapitulated critical features of cellular heterogeneity discovered in human GBM. The advantage that all LEGOs were derived from the exact iPSC clone with defined mutations allows us to further analyze how genetic heterogeneity contributes to cellular heterogeneity, which was not possible based on previous models. We first used t-SNE (t-distributed stochastic neighbor embedding) analysis for cell cluster analysis of all LEGOs together. There were 26 cell clusters we identified in one-month-old LEGOs (Figure 2I). Interestingly, we found that many cell clusters were dominated by cells from a single genotype (clusters 2, 5, 6, 7, 10, 13,17, and 22). In contrast, the other major clusters contain cells from multiple genotypes (clusters 1, 3, 4, 8, 9, 11, 12, 14, and 15) (Figure S2G), suggesting these cells are less dependent on cell genotypes. We then analyzed major lineage markers for stem cells and differentiation in the t-SNE plots. SOX2 positive stem cells were distributed to several cell clusters 1, 2, 7, 8, 10, and 17 (Figure 2J). Lineage marker expression showed that clusters 8 (mixed by PTN and WT) and 12 (mixed by PT and PTCC) are SOX2 positive stem cells (Figure 2J). Cluster 3 (mixed by PTN and WT) and 4 (mixed by PT and PTCC) were cells that express DCX (Figure 2J). APOE clusters are dominated mainly by cells of a single genotype (Figure 2J). The influence of genetic mutation on cell clusters is more pronounced in 4-month-old LEGOs; out of 34 clusters, most of the big clusters are dominated by cells from single genotypes (Figure 2I, S2H). Lineage marker expression on the t-SNE plot also suggests that similar lineage still stay close to each other on the t-SNE plots, but a clear difference was observed between different genotypes (Figure 2K). These results demonstrate that tumor mutations have a strong influence on cell phenotypes. However, the stem cell differentiation hierarchy governed by neurodevelopmental programs still operates during tumor formation.

### DNA methylome analysis reveals genotype-dependent progressive changes of DNA methylation during gliomagenesis

Tumor cell DNA methylation was recently used for the molecular classification of brain tumors ^6^. However, how different genetic mutations affect the DNA methylation pattern in GBM remains largely unclear. We selected the one-, two-, and three-month-old LEGOs and WT organoids for DNA methylation analysis using an EPIC (850K) DNA methylation array. A principle component analysis (PCA) suggests that the DNA methylome of WT organoids changes gradually over time, indicating a maturation signature of DNA methylation along PC2 (Figure 3A). Interestingly, the one-month-old PT and PTCC organoids are similar to the WT organoids (Figure 3A), indicating that these mutations do not lead to immediate dramatic DNA methylome changes. The PTN organoids differ from PT and PTCC already at one month of age (Figure 3A). Moreover, all LEGOs showed reduced progression along the maturation axis (PC2) compared to WT organoids (Figure 3A). This also indicates a sign of differentiation blockage, consistent with the scRNA-seq results. On the other hand, PC1 exhibits a gradual but genotype-specific change in DNA methylome (Figure 3A), suggesting oncogenic mutations induce genotype-specific DNA methylation changes. We then identified differentially methylated probes (DMP) among all groups at different developmental stages and found that DMP numbers were significantly different, with PT organoids showing the lowest and PTN organoids exhibiting the highest (Figure 3B). Interestingly, the methylation level of the mesenchymal subtype was also shown to be the highest among all three GB subtypes (Figure S3A) ^26^. The dynamic changes of DNA methylation in the LEGOs over time demonstrate that the DNA methylome is actively changing during tumor progression (Figure 3B), including both hypomethylated and hypermethylated probes, particularly during the early developmental stage of brain tumors (Figure 3C). However, it remains unclear what regulates these dynamic DNA methylome changes during tumor development. We performed a gene set enrichment analysis (GSEA) on the DMPs located on different gene features at various stages. There were no enriched hallmark gene sets in PT, probably due to the low number of DMPs. For the probes located 0-200 bp upstream of the transcription starting site in three-month-old PTCC organoids, we identified the enrichment of several hallmark gene sets, such as angiogenesis and interferon alpha response (Figure 3D). The most apparent difference was observed in the PTN group with strong activation of EMT and inflammatory signatures in different gene feature locations at three months and in the 5’UTR at two months (Figures 3D, 3E, S3B), in line with its infiltrative growth pattern *in vivo*.

**Figure 3.**
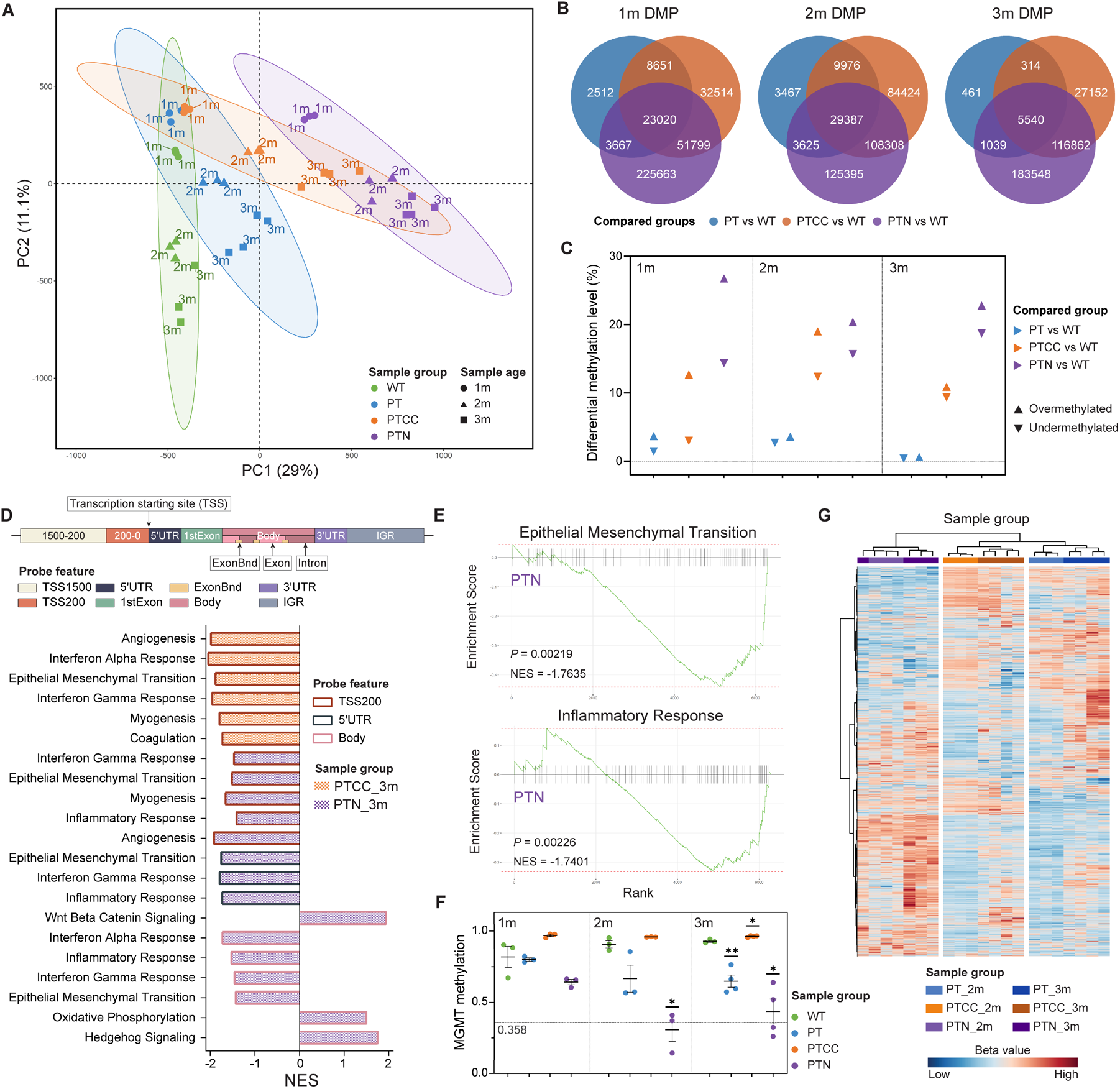
DNA methylome analysis reveals genotype-dependent progressive changes of DNA methylation during gliomagenesis. (A). PCA analysis of one-, two- and three-month-old organoids’ methylome, the ovals indicate 95% confidence ellipse of each genotype. (B). Venn diagrams demonstrate unique and common DMPs in LEGOs at one, two, and three months compared to age-matched WT. (C). Differential methylation level comparing LEGOs to age-matched WT control. (D). Probe gene feature distribution and GSEA hallmark enrichment of the DMPs located on different gene features (adjusted *P* value < 0.05) in three-month-old LEGOs. TSS, transcription starting site, UTR, untranslated region, IGR, intergenic region, ExonBnd, exon boundaries. (E). GSEA enrichment plots for the two hallmark gene sets enriched at the 5’UTR region in the three-month-old PTN organoids. (F). MGMT promoter methylation probability estimation in different organoids. N = 3 for all groups of one- and two-month-old organoids and three-month-old WT organoids, and N = 4 for three-month-old LEGOs. Data are represented as mean ± SEM. P values were calculated by Student’s t-tests comparing LEGOs to age-matched WT, and only significant values are labeled. ** *P* < 0.01, * *P* < 0.05. (G). Methylation classification heatmap with 8000 GB probes for two- and three-month-old LEGOs. See also Figure S3, Table S1

MGMT (O^6^-methylguanine-DNA methyltransferase) promoter methylation is associated with better TMZ response in GBM patients ^27^. Interestingly, we observed an increased level of MGMT promoter methylation in PTCC organoids compared to PT and PTN organoids (Figure 3F) which indicates that PTCC organoids may respond better to TMZ treatment than PT and PTN organoids. Moreover, unsupervised cluster analysis demonstrates that different LEGOs can be categorized by human GBM DNA methylation classification probes ^26^ (Figure 3G), indicating a human GBM-like methylation pattern in the LEGO model.

It has been suggested that DNA methylation signatures can be used to determine the cell of origin in human cancers ^28^. Our analysis demonstrated that the DNA methylome is dynamic during tumor development and is dependent on the mutation spectrum. Therefore, it is crucial to use stable and mutation-independent DNA methylation patterns as tracers for cancer cell origin. We generated a probe set (Table S1) that shows no significant changes among all different groups of organoids. Gene ontology (GO) analysis indicates that these probes are highly enriched for tissue development and differentiation (Figure S3C). This probe set can be further explored as candidates to trace brain tumor origins.

### Metabolic reprogramming and metabolic heterogeneity during brain tumor development

One of the hallmarks of cancer cells is the dysregulation of metabolism ^29^. However, it remains unclear how genetic heterogeneity affects the metabolic status of cancer cells. Therefore, we analyzed the intra- and extracellular metabolome of one- and four-month-old LEGOs and WT organoids (Figure 4A).

**Figure 4.**
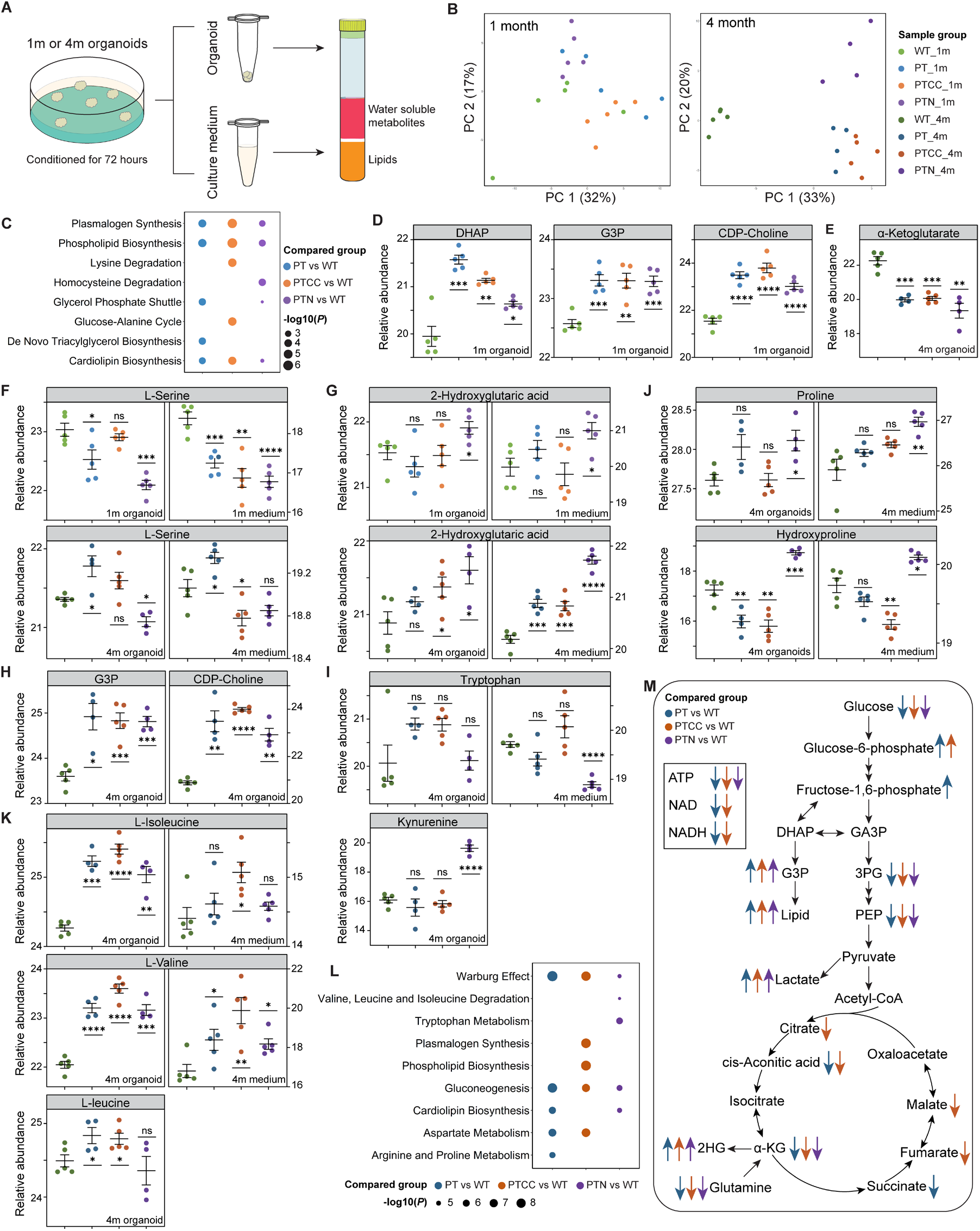
Metabolic reprogramming and metabolic heterogeneity during brain tumor development. (A). Schematic illustration of sample collection and extraction for metabolomic and lipidomic analysis. (B). PCA of one- and four-month-old organoid metabolome. (C). Dot plot shows the top five enriched pathways from the quantitative enrichment analysis of one-month-old LEGOs compared to WT organoids. (D). The relative abundance of lipid metabolism-related metabolites in one-month-old organoids. (E). The relative abundance of α-ketoglutarate in four-month-old organoids. (F). The relative abundance of L-serine in one- and four-month-old organoids and culture medium. (G). The relative abundance of 2-hydroxyglutaric acid in one- and four-month-old organoids and culture medium. (H). The relative abundance of lipid metabolism-related metabolites in four-month-old organoids. (I). The relative abundance of tryptophan (upper panel) in four-month-old organoids and culture medium and of the tryptophan metabolite kynurenine (lower panel) in four-month-old organoids. (J). The relative abundance of the major amino acids constituting collagen in four-month-old organoids and culture medium. (K). The relative abundance of branched-chain amino acids in four-month-old organoids and culture medium. (L). Top five enriched pathways from the quantitative enrichment analysis of the metabolites from four-month-old LEGOs compared to WT organoids. (M). Diagram demonstrating the metabolic changes in LEGOs. GA3P, glyceraldehyde-3-phosphate; 3PG, 3-phosphoglyceric acid; PEP, phosphoenolpyruvic acid. In D, E, F, G, H, I, J, and K, the color of the dots indicates the sample group, data are represented as mean ± SEM; N = 4 for four-month-old PT and PTN organoid samples, and N = 5 for the rest of the groups; *P* values were calculated with Student’s t-tests comparing LEGOs to WT; **** *P* < 0.0001, *** *P* < 0.001, ** *P* < 0.01, * *P* < 0.05, and ns, non-significant. See also Figure S4.

The metabolome of one-month-old organoids is largely similar to each other (Figures 4B, S4A, S4C). However, enrichment analysis of group-specific changes compared to WT organoids suggests the activation of phospholipid synthesis and glycerol phosphate shuttle in LEGOs (Figure 4C), indicated by the increase of DHAP (dihydroxyacetone phosphate), G3P (glycerol-3-phosphate) and CDP-choline (Figures 4D, S4A). This is consistent with our previous finding that GPD1 (glycerol-3-phosphate dehydrogenase 1), which converts DHAP into G3P, is specifically expressed in brain tumor stem cells but not in neural stem cells ^30^.

The four-month-old organoids’ metabolome showed a clear difference between LEGOs and WT organoids (Figure 4B). PT was similar to PTCC, while PTN was very distinct from the other LEGOs (Figure 4B). Heatmap analysis of intracellular metabolites demonstrates activation of the glycolysis pathway in LEGOs, indicated by low levels of glucose and glutamine and high levels of lactic acid, further confirmed by the medium metabolite data (Figure S4B, S4D, S4E). The TCA (tricarboxylic acid) cycle metabolites (citric acid, aconitic acid, α-ketoglutarate, succinate, fumarate, malate, ATP, NAD) were decreased in LEGOs compared to the WT organoids (Figure 4E, S4B, S4F), suggesting a shift toward glycolysis from oxidative phosphorylation, reminiscent of the Warburg effect.

Metabolites are essential substrates of many epigenetic enzymes ^31^. We analyzed metabolite changes that may explain the DNA methylome changes in the LEGOs. Serine contributes to methylation via the major methyl group donor S-adenosylmethionine ^32^. In one-month-old organoids, the level of serine in the culture medium was reduced in all LEGOs compared to WT organoids (Figure 4F, S4A, S4C), and the intracellular level of serine was most significantly decreased in the PTN organoids (Figure 4F). In contrast, in four-month-old LEGOs, the utilization of serine was increased in PTCC and PTN, while decreased in PT (Figure 4F, S4B, S4D). This strongly suggests that serine is consumed by all LEGOs and even more by the PTN organoids, which is in line with the observed high levels of hypermethylation in one- and three-month-old PTCC and PTN organoids (Figure 3C). Oxoglutaric acid (α-ketoglutarate, α-KG) is the substrate of many α-KG-dependent dioxygenases, including the DNA demethylation enzymes TET1/2/3 and 2-hydoxyglutaric acid (2-HG) antagonizes the function of α-KG ^31^. The level of 2-HG increased in one-month-old PTN extra- and intracellularly, and accumulated in four-month-old PTCC and PTN organoids as well as in all LEGO culture media, particularly in the PTN group (Figure 4G). In contrast, the level of α-KG was depleted in all four-month-old LEGOs compared to the WT organoids (Figure 4E). This further explains the dynamic DNA methylation changes in LEGOs and supports the hypermethylation pattern of PTN organoids.

Consistent with the results from one-month-old organoids, G3P and CDP-choline levels are higher in LEGOs at four months of age (Figure 4H), suggesting the mutant organoids depend on this lipid metabolism pathway. Regarding genotype-specific changes, we found that PTN organoids uniquely upregulate the tryptophan metabolism pathway by consuming and utilizing more tryptophan and producing more kynurenine (Figure 4I). Kynurenine could be catabolized into NAD to facilitate energy production, cellular proliferation, and immune suppression ^33,34^. PTN organoids also have high levels of proline and hydroxyproline in the organoids and culture medium (Figure 4J). Proline and hydroxyproline are the major amino acid components of collagen proteins ^35^. Collagen serves as the scaffold to facilitate glioma cell migration, increase the stiffness of the tumor, and induce an immune suppressive microenvironment ^36^,^37^, and elevated levels of hydroxyproline could be indicative of high collagen turnover. In PTCC organoids, we observed the accumulation of branched-chain amino acids (valine, isoleucine, and leucine) in both the organoids and the medium (Figure 4K), indicating an abnormal branched-chain amino acid metabolism. Enrichment analysis suggests that the Warburg effect is enriched in all LEGOs, with PTCC particularly showing enrichment of phospholipid biosynthesis, whereas tryptophan metabolism is among the most enriched pathways in PTN organoids (Figure 4L).

The results above demonstrate distinct metabolic reprogramming events during tumor development (Figure 4M), and it is evident that genetic mutations determine the metabolic differences in cancer cells. In addition, some metabolic changes may regulate the DNA methylome changes.

### Lipidomics assay uncovers glycerol lipid metabolism being a hallmark of GBM

The metabolomic analysis identified that the metabolites (DHAP, G3P, CDP-choline) in phospholipid biosynthesis are strongly associated with GBM development. We therefore performed lipidomic analysis using the same experimental setup shown in Figure 4A. The PCA analysis of one-month-old organoids shows that the lipidomes of LEGOs are different from WT organoids (Figure 5A). This was unlike the metabolome and methylome results, suggesting that lipidome reprogramming is the pioneering event upon the loss of tumor suppressors. The heatmap of lipid species indicates both DG (diacylglycerols) and TG (triacylglycerols) upregulation in all mutant groups (Figure 5B), which further illustrates the consequence of increased DHAP and G3P. This was further confirmed by enrichment analysis showing that TGs are the most significantly enriched lipid species in all LEGOs (Figure 5C). In addition, a decrease in ether-linked phosphatidylethanolamine (O-PE) was observed in the PTCC organoids (Figure 5B, 5C).

**Figure 5.**
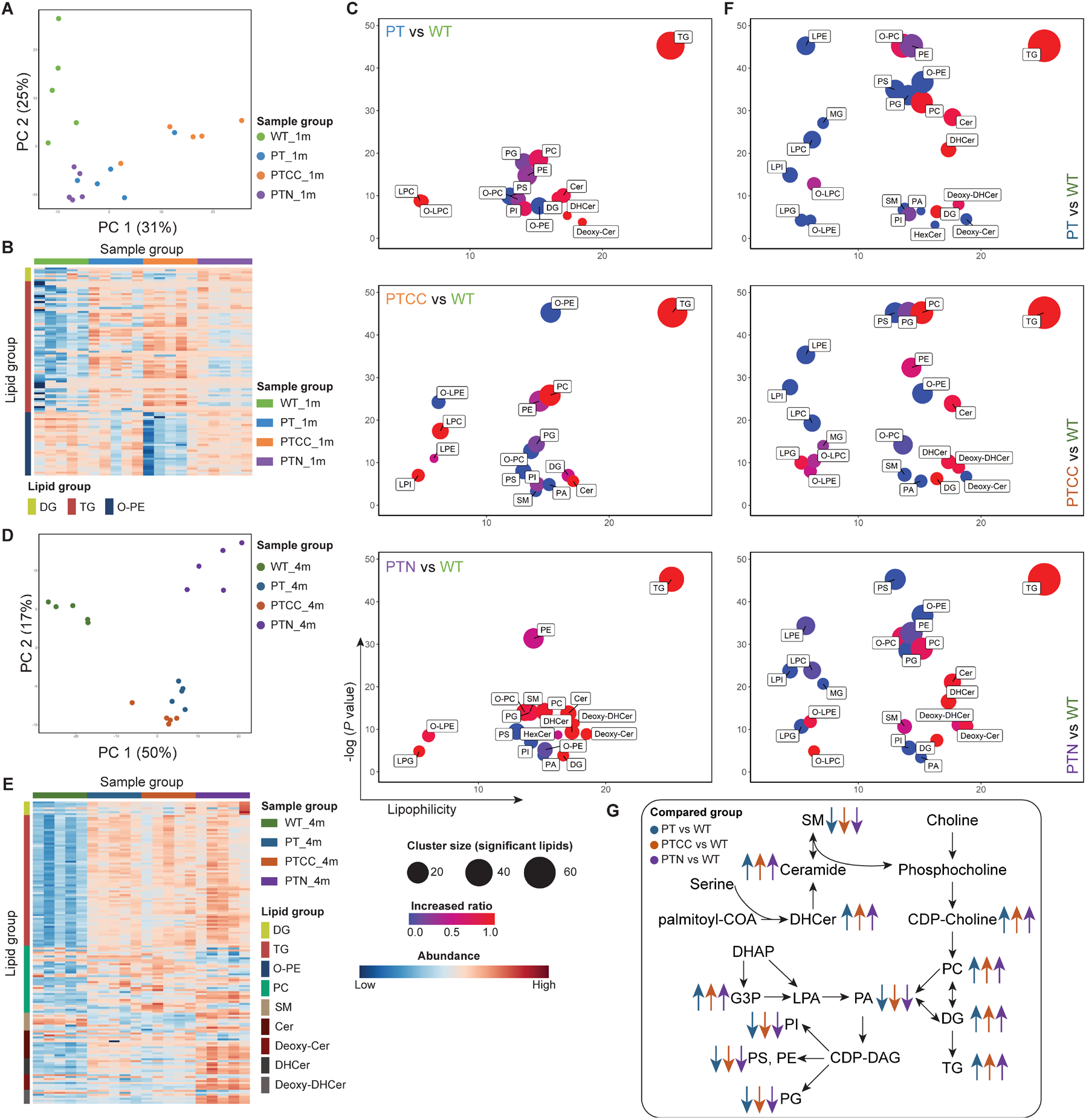
Lipidomics assay uncovers glycerol lipid metabolism being a hallmark of GBM. (A). PCA of the one-month-old organoid lipidome (B). Abundance heatmap of different lipids in one-month-old organoids. (C). Enrichment plot for different lipid groups in one-month-old organoids. (D). PCA of four-month-old organoid lipidome. (E). Abundance heatmap of different lipids in four-month-old organoids. (F). Enrichment plot for different lipid groups in four-month-old organoids. (G). Diagram demonstrating the lipidomic changes in the LEGOs compared to WT organoids. In C and F, the cluster size indicates the number of significantly changed lipids in the group comparison. The increased ratio was calculated by dividing the number of significantly increased lipids by the total number of significantly changed lipids. See also Figure S5.

We next analyzed the lipidome of four-month-old organoids. PCA was similar to the four-month-old metabolome PCA, with the leading principal component (PC1) separating the LEGOs from WT and the second principal component (PC2) distinguishing PTN from PT and PTCC (Figure 5D). DGs, TGs, and phosphatidylcholine (PC) were significantly increased in all mutant organoids, particularly in PTN (Figure 5E, 5F). This, together with the increase of G3P, DHAP, and CDP-choline, as described above, highlights the importance of TG and choline metabolism in GBM (Figure 5G). On the other hand, the structural phospholipids (such as PG, PI, PS, PE, and O-PE) are decreased in mutant organoids (Figure S5A). It is likely that the increased production of DG, TG, and PC in LEGOs leads to decreased structural phospholipids as these lipids are derived from the same precursor, G3P. Ceramide production could be activated under stress conditions by hydrolyzing sphingomyelin (SM) ^38^. Consistently, we observed a higher amount of SM in WT, and abundant ceramides and CDP-choline in all LEGO groups (Figure 5E, 5F, 5G, 4H), suggesting augmented activation of SM hydrolysis. PTN exhibited significantly higher ceramide expression than all other groups (Figure 5E), implying a unique mechanism enhancing ceramide synthesis upon loss of *NF1.* It was shown that the tryptophan metabolite kynurenine can directly bind and activate the aryl hydrocarbon receptor (AHR) ^34,39^, and the activation of AHR elevates the synthesis of ceramides ^40,41^. Altogether, the lipidome analysis identified that lipid reprogramming is a pioneering event during gliomagenesis, and glycerol lipid metabolism is a hallmark of GBM (Figure 5G).

### Proteomic/phospho-proteomic analysis identifies actionable targets and pathways for the genotype-based treatment of GBM

To search for possible genotype-specific drug targets using the LEGO models, we next performed proteomic and phospho-proteomic analyses on four-month-old organoids. Proteome and phospho-proteome PCA plots exhibit high similarity to the metabolome and lipidome, with a distinct difference between LEGOs and WT organoids, and PTN shows a more distinct proteome/phospho-proteome profile compared to PT and PTCC (Figure 6A). However, phospho-proteome provided a better separation between PT and PTCC (Figure 6A). GSEA analysis of differentially expressed proteins identifies processes involved in LEGO development (Table S2). In particular, the cholesterol and lipid pathways are enriched in all LEGOs (Figures 6B, 6C, 6D), consistent with the metabolomic and lipidomic data. The G2M checkpoint and related stress and mitosis pathways are enriched in PTCC (Figure 6C), indicating elevated mitosis as a result of *CDKN2A/2B* deletion. PTN organoids are enriched for negative regulation of immune response and SASP (senescence-associated secretory phenotypes) (Figure 6D, S6A). Additionally, signatures associated with DNA methylation, extracellular matrix disassembly, as well as several collagen proteins are highly enriched in PTN (Figures 6D, S6B). This is concordant with the observed DNA methylome changes, the *in vivo* infiltrative phenotype of PTN tumors, and the proline/hydroxyproline enrichment in PTN metabolome, respectively.

**Figure 6.**
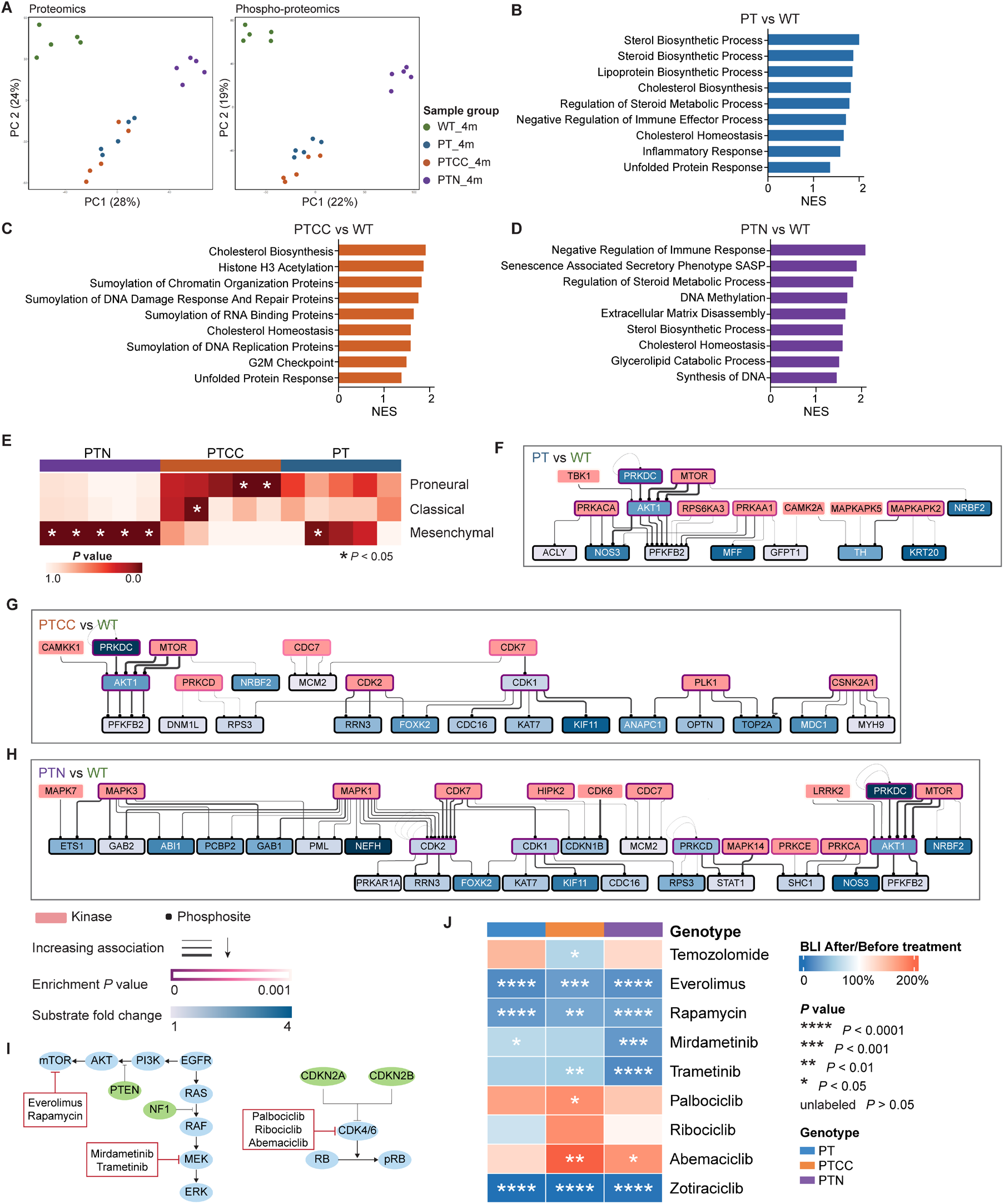
Proteomic/phospho-proteomic analysis identifies actionable targets and pathways for the genotype-based treatment of GBM. (A). PCA of proteomics and phospho-proteomics of four-month-old organoids. Representative significantly enriched pathways or terms in PT (B), PTCC (C), and PTN (D) compared to WT identified by GSEA. (E). Heatmap showing the enrichment of GBM subtype signatures ^42^ in different LEGOs. Enriched kinases together with their substrates in PT (F), PTCC (G), and PTN (H) compared to WT. (I). Illustration of selected drugs and their targets. (J). Treatment outcome for the proof-of-principle drug tests. N = 3 for TMZ treatment groups, N = 4 for kinase inhibitor treatments. *P* values were calculated with paired Student’s t-tests comparing signals measured after treatments to before treatments See also Figure S6, Table S2, Table S3, Table S4

RNA expression has been used for the molecular classification of GBM, but little is known about whether using protein expression as the classifier will yield similar results. We analyzed our proteome data using established tumor-cell-specific RNA signatures ^42^ and found that mesenchymal signatures are highly enriched in the PTN organoids (Figure 6E, S6C). This result, together with the other omics analyses, firmly confirms that the PTN organoid resembles the mesenchymal subtype of GBM. Furthermore, we found that the expression of MGMT protein is high in the PTN group, and expression of IDH1 is increased in PT and PTCC compared to WT, concordant with the methylation changes (Figure S6D).

To identify possible actionable targets in different subgroups of LEGOs, we utilized a drug-gene interaction database ^43^ to identify druggable targets for LEGOs (Table S3). Collectively, enzymes involved in lipid metabolism enzymes, such as MGLL (monoglyceride lipase), and FDFT1 (farnesyl-diphosphate farnesyltransferase 1), could be potential targets for all LEGOs (Figure S6E); this is in line with the activation of lipid metabolism in LEGOs. DNMT3A (DNA methyltransferase 3A), SPTLC2 (serine palmitoyltransferase 2), and cholinesterase (BCHE) could be interesting targets for PTN (Figure S6E).

The phospho-proteomic data allow the prediction of possible kinases involved in tumor progression. We used a kinase-target interaction database ^44^ and Kinase Enrichment Analysis ^45^ to identify the upstream kinases of the phosphorylated sites (Table S4). PT organoids showed activation of AKT1 and mTOR, due to the mutation of *PTEN* (Figure 6F). Surprisingly, the PTCC organoid phospho-proteomic data did not show enrichment of CDK4/6, which are classic substrate kinases of CDKN2A/2B. Instead, CDK1/2/7 were activated in addition to mTOR and AKT1 (Figure 6G). In addition to AKT1 and mTOR, MAPK1, MAPK3 and CDK7 were upregulated due to *NF1* mutation in PTN organoids (Figure 6H). Using luciferase as a readout for tumor cells in the LEGO model, we found that PTCC LEGO is more sensitive than the other two to TMZ (Figure 6J), in line with a higher MGMT promoter methylation status. the mTOR inhibitors were effective in all LEGOs (Figure 6I, 6J, S6F, S6G), consistent with the kinase enrichment analysis. In contrast, CDK4/6 inhibitors exhibited no growth inhibition in all LEGOs (Figure 6J), again concordant with the kinase enrichment analysis, which indicated that CDK4/6 were either not enriched or only showed low enrichment compared to other CDKs. On the other hand, MEK1/2 inhibitors were effective in all groups but most effective in PTN organoids, likely because of the enhanced activation of MAPKs (Figure 6J). Combination therapy using mTOR and MEK inhibitors show the most effective inhibition of PT and PTN growth, while CDK4/6 inhibitors compromise mTOR inhibitor effects in PTCC organoids (Figure S6H), suggesting that CDK4/6 inhibitors should be carefully examined before being considered for treating GBM patients with *CDKN2A/2B* mutations. Moreover, we treated the LEGOs with the CDK inhibitor Zotiraciclib targeting CDK1/2, which was highly activated in all the mutant organoids, and observed that all LEGOs were highly sensitive to Zotiraciclib treatment (Figure 6J), suggesting that CDK1/2 are valuable therapeutic targets in GBM.

### LEGO-based drug screening identifies novel drug candidates for GBM therapy

With the goal of generating a genotype-based drug reference map and possibly identifying new treatment strategies for GBM, we performed a drug screen on 327 drugs containing FDA-approved drugs that could penetrate through the blood-brain barrier (Figure. 7A, Table S5). All LEGO cells were engineered to express luciferase, and the bioluminescence signal was used as a readout of cell numbers in the LEGOs. To select effective drugs, among the drugs that resulted in significant inhibition of bioluminescence signal (*P* < 0.05), we only selected drugs that resulted in 50% inhibition of bioluminescence signal as positive candidates. With this screen, we identified 42 drugs with therapeutic effects; seven acted on all three genotypes, and the rest only worked on specific genotypes (Figure 7B, 7C, 7D). We found that EGFR inhibitors Dacomitinib and Osimertinib inhibit LEGO growth in all genotypes, with a particularly strong effect on the PTN organoid (Figure 7C, 7D, 7E), suggesting a strategy of patient enrollment for clinical trials for testing EGFR inhibitors. The Syk (spleen tyrosine kinase) inhibitor Fostamatinib inhibits all LEGOs, suggesting that Syk signaling is essential for GBM progression (Figure 7C). Interestingly, we also found that the schizophrenia drug Aripiprazole also inhibits tumor growth in all LEGOs (Figure 7C, 7F), which implies an alteration of dopamine signaling in GBM.

**Figure 7.**
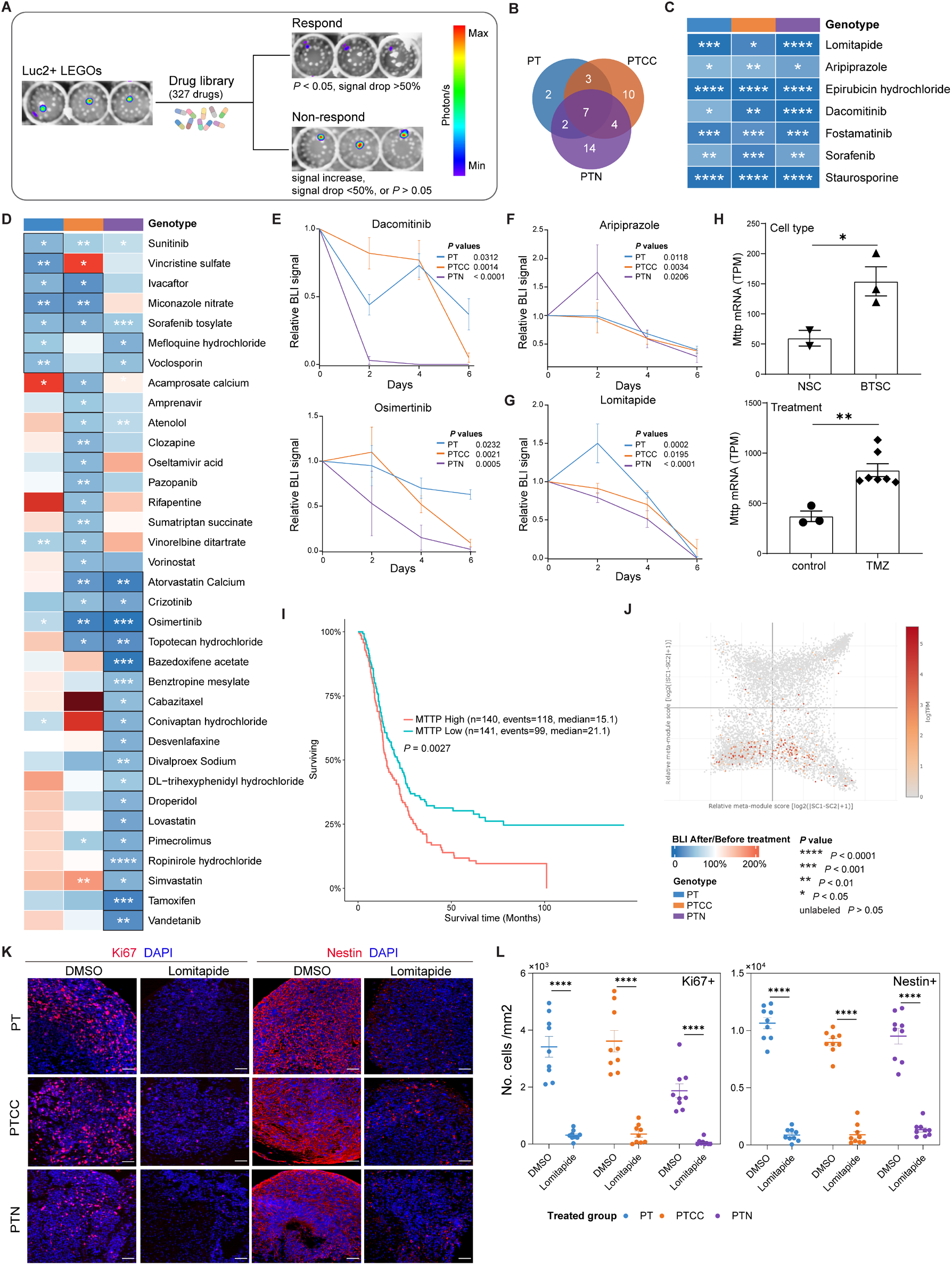
LEGO organoids respond to drugs that target mutation-specific mechanisms. (A). Illustration of BLI-based drug screen. (B). Venn diagrams demonstrating effective drug distribution in different LEGOs (C). Treatment outcome for drugs effective in all LEGOs. N = 3 for each group. (D). Treatment outcome for drugs that are effective in at least one group. The black frames around the cells highlight the effective group. N = 3 for each group. Cell viability tracing with BLI signal in LEGOs treated with Dacomitinib and Osimertinib (E), Aripiprazole (F), and Lomitapide (G). (H). Normalized Mttp expression from ribo-seq analysis of mouse BTSC and NSC (left) and RNA-seq analysis of mouse brain tumors treated with/without TMZ (right) (I). MTTP expression on GBM patient survival from an external data set ^46^, *P* value was calculated with Log rank test (J). MTTP expression from GBM patient single-cell RNA sequencing data. (K). Representative Ki67 and Nestin staining on LEGOs treated with DMSO or Lomitapide. Scale bar, 50 μm. (L). Quantification of Ki67^+^ and Nestin^+^ cells treated with lomitapide, N = 9 sections for each group. In C and D, *P* values were calculated with paired Student’s t-tests comparing signals measured after treatments to before treatments; **** *P* < 0.0001, *** *P* < 0.001, ** *P* < 0.01, * *P* < 0.05. In E, F and G, data are represented as mean ± SEM. See also Table S5

Most interestingly, we found that Lomitapide, an inhibitor of microsomal triglyceride-transfer protein (MTTP), inhibits tumor growth in all LEGOs (Figure 7C, 7G). MTTP is a lipid transfer protein and is essential for the regulation of lipid metabolism. This is in line with our discovery that glycerol lipid metabolism is a hallmark of GBM metabolism. Expression of Mttp has highly enriched in our previous ribosome RNA-sequencing analysis by comparing mouse neural stem cells (NSCs) and brain tumor stem cells (BTSCs), its expression is also highly enriched in tumor-bearing mice after TMZ treatment (Figure 7H) ^30^. High expression of MTTP also shows a worse prognosis in GBM patient (Figure 7I) ^46^, and single-cell analysis in published data sets ^9^ suggest MTTP is more expressed in stem cell or mesenchymal subtype of tumor cells (Figure 7J), which is consistent with our previous observation of activation of glycerol metabolism in BTSCs ^30^. In the Lomitapide treated LEGOs, we found a striking reduction in the number of proliferation cells and stem cells (Figure 7K, 7L). Surprisingly, ER (estrogen receptor) modulator Tamoxifen and Bazedoxifene acetate exhibited selection inhibition of PTN organoid, warranting additional investigation of ER function in mesenchymal GBM (Figure 7D).

Complete information on treated drugs and outcomes can be found in Supplementary Table 5. Noteworthily, some drugs that exhibited therapeutic effects on one genotype may promote the growth of another, which further highlights the importance of genetic background in directing treatment options. This genotype-based drug reference provides a basis for the personalized treatment of GBM patients.

## Discussion

Our temporal multi-omics analysis (scRNA-seq, DNA methylome, Metabolome/Lipidome, and Proteome/Phospho-proteome) covers essential molecular layers of the cancer cell molecular network. This allows us to discover genotype-specific molecular changes during tumor development. In Table S6 (supplementary information), we summarized all major molecular milestones during GBM development and divided them into shared and genotype-specific milestones. We also list milestones that can be validated by analysis of different molecular layers. i.e., early changes during GBM development include the increase of stem cell frequency and attenuation of neural differentiation. This is accompanied by metabolite changes, which can also influence epigenetic modifications like DNA methylation. Coherently, active DNA methylation changes during early tumor development shown by methylation array data, elevated DNA methylation pathway activity, and low IDH1 expression presented by proteomics data support hypermethylation in PTN, which could be further confirmed by the decrease of a-KG and increase of 2-HG in the metabolomic assay. The increase of phospholipid metabolism is an early event, and this change persists with brain tumor development. Notable genotype-specific features include a WNT activation in PTCC organoids, ectopic expression of the HOX gene cluster, and mesenchymal signature in PTN organoids. The MGMT promoter is methylated in PTCC organoids, and we show that PTCC organoids are sensitive to TMZ. It is also important to note that the LEGOs are primarily similar to the WT organoids at the one-month-old, highlighting that most of the oncogenic changes occurred during tumor organoid development, not at the iPSC stage.

One fundamental question in cancer biology is which features of cancer cells are determined by genetic and non-genetic heterogeneity, respectively. This could not be investigated so far due to the lack of proper models. The genetically defined LEGOs are initially derived from the exact iPSC clone providing an ideal tool to assess the contribution of genetic heterogeneity to intratumoral heterogeneity quantitatively. In our analysis, *CDKN2A/2B* mutation in *PTEN* and *TP53* deletion background further push the development of PT organoids in a similar direction, suggesting these mutations work together and drive similar cancer phenotypes. However, the *NF1* mutation dramatically reprograms the cancer cell phenotypes across all molecular layers, which will be discussed below. Therefore, the LEGO model can serve as genetic building blocks of the cancer genome which can be further expanded and used to analyze the interaction between cancer genetic and non-genetic heterogeneity. Combining LEGOs to generate fully customized genetically heterogenous organoids is also straightforward. The scRNA results also demonstrated that genetic mutations have mutation specific influences on cell phenotypes. Although the stem cell differentiation hierarchy is largely maintained in all LEGOS. The cellular composition and molecular phenotype of the lineages in different LEGOs are different from each other. This is critical information for future interpretation of scRNA-seq results of human GBM patient tissues, the contribution to cellular heterogeneity from genetic and non-genetic factors must be clearly demonstrated. Therefore, obtaining mutation information and considering the genetic heterogeneity within different cell clusters is essential before claiming they may represent different cell states ^9^.

Another striking observation in our multi-omics analysis is the activation of phospholipid metabolism throughout LEGO development. Interestingly, this activation is already noticeable in one-month-old LEGOs, supported by increased DHAP and G3P. DHAP is the intermediate metabolite of glycolysis and can be converted by GPD1 into G3P, the primary precursor for lipid metabolism. We have shown before that GPD1 is induced explicitly in brain tumor stem cells during brain tumor development and blocking GPD1 alters tumor lipid metabolism and prolongs the survival of brain tumor-bearing animals ^30^. The increase of DHAP, G3P, and CDP-choline in the metabolomic analysis and the increase of DG, TG, and PC in the lipidomic analysis in LEGOs demonstrate that lipid metabolism, particularly the glycerophospholipid metabolism, is activated during brain tumor development. This is in line with the clinical observation that the glycerol level in GBM patients is much higher in tumors compared to normal tissue in the tumor periphery ^47^. It was also reported that brain metastasis also upregulates lipid metabolism ^48,49^, indicating an adaptation of cancer cells to the lipid-deprived brain environment ^48,50^. For this purpose, brain tumor cells upregulate GPD1 to switch the metabolic flow to lipid metabolism by making use of the glycolysis metabolite DHAP, which was also reported to be the only sensor metabolite of the mTOR pathway in glycolysis ^51^. More importantly, we also discovered that MTTP inhibitor Lomitapide efficiently blocks LEGO growth, providing another attractive target, and the drug should be further investigated. Lipid metabolism is likely an emerging hallmark of brain cancers that should be further investigated.

Major mutations that drive human GBM have been identified via genomic sequencing ^4,5^. Interestingly, the major molecular subtypes of human GBM are defined primarily via RNA expression or DNA methylation pattern ^8,42,52^, and there is no strong correlation between genetic mutation and molecular subtypes. *NF1* mutation is highly enriched in the mesenchymal subtype, whereas *TP53*, *PTEN*, and *CDKN2A/2B* inactivation were not enriched in particular subtypes ^4^. Inactivation of *Nf1* and *Tp53* leads to brain tumor formation in a mouse model ^53^. However, whether *NF1* mutation drives mesenchymal GBM remains not experimentally confirmed. Here we showed that *NF1* mutant organoids have many unique features compared to other LEGOs. The PTN xenograft shows a rather infiltrative growth pattern and high angiogenesis, and scRNA-seq identified a mesenchymal cell cluster with increased expression of collagen genes. PTN also produces the immunosuppressant kynurenine and has high levels of proline and hydroxyproline, which support the high collagen level. Moreover, PTNs are not sensitive to TMZ treatment because of lacking MGMT methylation and increased expression of MGMT protein. All these factors fit the mesenchymal features of human GBM ^23^ and confirm that *NF1* mutation drives the mesenchymal features in human GBM. The significant differences between PTN tumors and PT/PTCC tumors suggest that the *NF1* mutant GBM is a unique subgroup of GBM, which should be studied and treated differently.

The LEGO model analysis demonstrates that genetic mutations determine major molecular consequences. Therefore, the realization of personalized treatment of human GBM requires knowledge of genotype-specific drug sensitivity information. Our preliminary treatment of LEGOs demonstrates that different LEGOs respond differently to drug treatments. This set the foundation for using LEGO-like models to study human cancer heterogeneity. The results obtained from the LEGOs show an excellent correlation across different molecular layers, including drug responses. The MGMT promoter was found to be highly methylated in PTCC organoids, and the PTCC organoids respond better to TMZ treatment. In particular, it is unexpected that the PTCC organoid do not respond to CDK4/6 inhibitors and our phospho-proteome results suggest CDK4/6 are not activated in the PTCC organoids. This raises concern about using a *CDKN2A/2B* mutation as a selection criterion for CDK4/6 inhibitors. The drug screen we performed also provided precious information on a genotype-based drug sensitivity map, which can be used for drug candidate selections on personalized treatment GBM clinical trials. The following steps will further expand the LEGO genotypes and assemble different LEGOs to build fully customized, genetically heterogenous organoids, that can be used to investigate clonal evolution, cell competition, clonal interactions, and combination therapies. Moreover, adding tumor stromal cells like microglia and T cells will also be interesting, as it will allow investigations into how genetic heterogeneity determines immune cell behavior.

## Materials and methods

### Genome editing of iPSCs

Human induced pluripotent stem cells (iPSC) with mEGFP inserted at the safe harbor locus AAVS1 under CAGGS promoter were purchased from Coriell Institute (New Jersey, USA, Cat#AICS-0036-006; RRID: CVCL_JM19). All the iPSCs were cultured in Matrigel (Corning, New York, USA, Cat#354277) coated plates, fed with mTeSR Plus medium (Stemcell Technologies, Vancouver, Canada, Cat#100-0276) every other day at 37 °C incubators supplied with 5% CO_2_. The cells were passaged with ReleSR (Stemcell Technologies, Cat#05872) as small colonies after reaching 70% - 80% confluency. 3 μM of CHIR99021 (Tocris Bioscience, Minneapolis, USA, Cat#4423) and 1 μM of PD0325901 (Selleckchem, Houston, USA, Cat#S1036) ^19^ were added to the culture medium of PTN iPSCs. The cultures were regularly tested for mycoplasma contamination.

The gRNAs targeting respective tumor suppressor genes were inserted into modified pX330 plasmids ^54^ containing the puromycin-resistant gene. The electroporation was conducted with Neon^™^ Transfection System (Thermo Fisher Scientific, Massachusetts, USA). Briefly, cells were harvested by four minutes of Accutase (Sigma-Aldrich, Missouri, USA) treatment at 37 °C and resuspended with R resuspension buffer containing 15 μg gRNA expression vectors. The electroporation was conducted for two pulses with 1200 V, 20 ms. The electroporated cells were cultured in mTeSR Plus medium containing ROCK inhibitor (Stemcell Technologies, Cat#72304) for 24 hours after the electroporation. Puromycin (2 μg/mL) was added to the culture medium and refreshed every 12 hours for two days. The cells were then seeded at a density of 50 ~ 100 cells per 10 cm dish and expanded for ten days. The single cell colonies were screened by T7 Endonuclease I (T7E1, New England Biolabs, Ipswich, USA, Cat#M0302L) assay, and the Sanger sequencing (Eurofins) results were analyzed with the TIDE webtool ^55^. The A-tailing PCR products from the candidate clones were cloned into the pGEM-T vector (Promega, Madison, USA, Cat#A3600) and sequenced. Then two clones from each mutation combination were selected and further validated by western blotting. In brief, the protein lysis from the iPSCs was electrophoresed and transferred onto 0.2 μm PVDF membranes. The membranes were blocked with 5% non-fat milk for one hour at room temperature (RT) and incubated with primary antibodies overnight at 4 °C with shaking. The next day, the membranes were incubated with respective horseradish peroxidase (HRP) conjugated secondary antibodies for two hours and imaged using ChemiDoc (Bio-Rad, California, USA) after reacting with HRP substrate. gRNA sequences (5’ to 3’): *PTEN*, CAGTTTGTGGTCTGCCAGCT, *TP53*, GCAGTCACAGCACATGACGG, *CDKN2A*, GATGATGGGCAGCGCCCGAG, *CDKN2B*, CTGGCCAGCGCCGCGGCGCG, *NF1*, CCAGGATATATCCAAAGACG. Antibody dilutions: PTEN (Cell Signaling Technology, Massachusetts, USA, Cat#9559L, RRID: AB_390810), 1:1000; P53 (Thermo Fisher, Cat#MA512557, AB_10989883), 1:1000; P15/P16 (Santa Cruz, Texas, USA, Cat#sc-377412), 1:50; NF1 (DKFZ, Heidelberg, Germany, Cat#DKFZ-NF1-146/29/25 ^56^), 1:4; GAPDH (Cell Signaling Technology, Cat#2118L, Cat#2118L), 1:2000; β-Tubulin (Cell Signaling Technology, Cat#2128s, Cat#2128s), 1:2000.

### Organoid culture

The organoids were generated following previously described protocols ^20,57,58^ with minor adaptions. Briefly, on day 0, the iPSCs were dissociated into single cells as described above and seeded in 96-well ultra-low attachment plates (Corning, Cat#7007) containing the following medium, 80% DMEM/F12 (v/v, Gibco, Montana, USA, Cat#11330032), 20% KOSR (v/v, Gibco, Cat#10828-028), 3% ES-qualified fetal bovine serum (v/v, FBS, Gibco, Cat#10270106), 1% GlutaMAX (v/v, Gibco, Cat#35050038), 1% MEM-NEAA (v/v, Sigma-Aldrich, Cat#11140050) and 0.7% 2-Mercaptoethanol (v/v) supplied with 50 μM of ROCK inhibitor and 6 ng/mL of bFGF (Peprotech, New Jersey, USA, Cat#100-18B). The medium was refreshed on day 3. On day 5 the culture medium was replaced with DMEM/F12 containing 1% N2 (v/v, Gibco, Cat#17502048), 1% GlutaMAX (v/v), 1% MEM-NEAA (v/v), and 1 μg/mL heparin (Sigma-Aldrich, Cat#H3149) and cultured for four days. The EBs were then embedded in Matrigel and cultured in the following medium, 50% DMEM/F12 (v/v), 50% Neurobasal (v/v, Gibco, Cat#21103049), 0.5% N2 (v/v), 2% B27 without Vitamin A (v/v, Gibco, Cat#12587010), 0.025%insulin (v/v, Sigma-Aldrich, Cat#I9278), 0.35% 2-Mercaptoethanol (v/v,), 1% GlutaMAX (v/v), 0.5% MEM-NEAA (v/v), and 1% Penicillin/Streptomycin (v/v) for four additional days in 6 well ultra-low attachment plates (Corning, Cat#3473). Thereafter, the organoids were cultured on orbital shakers in culture medium containing 50% DMEM/F12 (v/v), 50% Neurobasal (v/v), 0.5% N2 (v/v), 2% B27 (v/v, Gibco, Cat#17504044), 0.025% insulin (v/v), 0.35% 2-Mercaptoethanol (v/v), 1% GlutaMAX (v/v), 0.5% MEM-NEAA (v/v), 1% Antibiotic-Antimycotic (v/v, Gibco, Cat#15240096) and 0.4 mM L-Ascorbic Acid (Sigma-Aldrich, Cat#A4544). The medium was exchanged every two to three days until sample collection. 3 μM of CHIR99021 and 1 μM of PD0325901 ^19^ were added to the PTN culture for the first 5 days. Widefield images for the organoids were taken by the Cell Observer (Zeiss, California, USA).

### Luciferase labeling of iPSCs

HEK293T cells were cultured with IMDM (Gibco, Cat#31980030) supplied with 10% FBS (v/v, ATCC, Cat#30-2020) at 37 °C incubators supplied with 5% CO_2_ and passaged with Trypsin/EDTA (Gibco, Cat#15400054). For lentivirus production, 5 × 10^6^ cells were seeded in 10 cm dishes and co-transfected with 2 μg envelope plasmid pMD2.G, 2 μg packaging plasmid psPAX2, and 4 μg luciferase (Luc2) expressing vector pHHLVX-EF1α-Luc2-puro the next day. DNA vectors were mixed with OptiMEM (Gibco, Cat#31985062) to a total volume of 250 μL, and 3x DNA volume polyethyleneimine (PEI, 1 mg/mL) was diluted in OptiMEM to a total volume of 250 μL, respectively. The two mixtures were then combined, thoroughly mixed, and incubated at RT for 15 minutes before being dropwise applied to the HEK293T cells. The virus particles were collected 24 hours and 48 hours after transfection and concentrated with Lenti-X^™^ Concentrator (Takara, California, USA, Cat#631232). The pellets were then resuspended with PBS, aliquoted, and stored at −80 °C. The iPSCs were infected with the lentivirus, and positive clones were selected by bioluminescence imaging (BLI) with IVIS Lumina II *In Vivo* Imaging system (PerkinElmer, Waltham, USA). Ten Luc2 positive clones were pooled to maximize the labeling rate and minimize the colony effect.

### Sample collection and cryosection

The organoids were fixed in 4% PFA for 20 minutes to 1 hour at 4 °C and emerged in 30% sucrose (w/v) overnight at 4 °C to dehydrate the tissue. The next day, the organoids were embedded in Gelatin/Sucrose solution and froze on dry ice. Gelatin/Sucrose solution was prepared by dissolving 7.5% gelatin (w/v) in 10% sucrose (w/v) at 37 °C. The embedded samples were stored in sealed plastic bags in a −80 °C freezer. Sections were collected and dried for one hour at RT before storing at −80 °C.

### Hematoxylin & Eosin (H&E) staining

The paraffin sections were deparaffined with the following procedure: 2× 5 minutes Xylene, 2×5 minutes 100% Ethanol, 2× 5 minutes 95% Ethanol, and 5 minutes 70% Ethanol. The sections (cryosections or deparaffined sections) were rehydrated in ddH_2_O for 5 minutes, stained in Hematoxylin for 1.5 minutes, and rinsed for 5 minutes under running tape water. 0.1% Eosin was applied for 1.5 minutes, washed by dipping in water, and differentiated in 70% Ethanol for 3 minutes. Dehydration was done with the following changing of buffers: 3 minutes 85% Ethanol, 2× 5 minutes 100% Ethanol, 2× 5 minutes Xylene. The sections were mounted with Eukitt and imaged with Axioscan (Zeiss) or Tissue FAXS Plus (Tissue Gnostics, California, USA).

### Immunofluorescence (IF) and Immunohistochemistry (IHC)

For IF staining, the sections were incubated with primary antibodies overnight at 4 °C after deparaffinization (for paraffin sections only), rehydration, antigen retrieval, and blocking. The sections were then incubated with respective secondary antibodies conjugated with AlexaFluor (Thermo Fisher Scientific, AF488, Cat#A11039, RRID: AB_2534096; AF555, Cat#A31572, RRID: AB_162543) or CF^®^ (633, Sigma-Aldrich, Cat#SAB4600128) dyes and DAPI for two hours at RT in the dark. The slides were mounted with Prolong gold (Invitrogen, California, USA, Cat# P36930) and imaged with Axioscan or Tissue FAXS Plus.

For IHC staining, the sections were first deparaffinized, rehydrated and antigen retrieved. Then they were treated with 3% H_2_O_2_ (v/v) for 10 minutes to quench the endogenous peroxidase before blocking and primary antibody incubation. On the second day, the sections were incubated with HRP conjugated secondary antibodies for two hours at RT. Then the sections were treated with Streptavidin HRP for 10 minutes and visualized by DAB substrate application (Abcam, Waltham, USA, Cat#ab64238). The cell nuclei were counterstained with Hematoxylin, then dehydrated and mounted as the H&E staining. For BrdU staining, the sections were treated with 2N HCl for 5 minutes at 37 °C before blocking. The slides were imaged with Axioscan or Tissue FAXS Plus.

Primary antibody dilutions: Ki67 (Cell Signaling Technology, Cat#9129, RRID: AB_823664), 1: 400 for IF, 1:1000 for IHC; GFP (Abcam, Cat#ab13970, RRID: AB_300798), 1: 500 for IF; GFAP (Cell Signaling Technology, Cat#3670, RRID: AB_561049), 1: 1000 for IHC; BrdU (BD Biosciences, New Jersey, USA, Cat#347580, RRID: AB_10015219), 1: 500 for IHC; Nestin (Cell Signaling Technology, Cat#33475, RRID: AB_10015219), 1: 200 for IF, 1:1000 for IHC; SOX2 (Abcam, Cat#ab97959, RRID: AB_2341193), IF 1: 500; Tubulin B3 (TUJ1) (Biolegend, California, USA, Cat#801202, RRID: AB_10063408), IF 1: 1000.

### Mouse orthotopic xenograft

Female NOD/SCID mice were purchased from Shanghai Jihui Laboratory Animal Care Co.,Ltd (Shanghai, China) and housed in the Animal Facility at the National Facility for Protein Science in Shanghai. All mouse experiments were conducted under Shanghai Institutional Animal Care and Use Committee (IACUC) guidelines and an approved IACUC protocol of ShanghaiTech University (#20201208001).

Single cells dissociated from the 45-day-old mutant organoids induced from Luc2+ iPSCs (PT, PTCC, PTN) were orthotopically injected into the right striatum of 4- to 5- week-old female NOD/SCID mice. Briefly, organoids were cut into small pieces with scalpels and digested with the Neural dissociation kit (P) (Miltenyi, Bergisch Gladbach, Germany, Cat#130-092-628) following the manufacturer’s protocol. 5 × 10^5^ cells were resuspended in 2 μL HBSS (Gibco, Cat# 14170088) and stored on ice. The mice were anesthetized with 0.15 ‰ tribromoethanol (Sigma-Aldrich, Cat#T48402), tribromoethanol (Sigma-Aldrich, Cat#T48402), and the cells were injected at the position 2 mm to the right lateral bregma and 3 mm deep with a flow of 0.2 μL/min utilizing a 10 μL precision micro syringe (World Precision Instruments, Florida, USA) with a 34-gauge needle. Mice were checked daily for signs of distress, including continuous weight loss or neurological disorders (such as hydrocephalus or impaired motor skills), and sacrificed with CO_2_ as soon as they showed related symptoms. The brains were collected for histological analysis.

### Single-cell RNA sequencing and data analysis

The single-cell RNA sequencing libraries were generated from one- and four-month-old whole organoid dissociations using Chromium Single Cell 3’ Kit v3.1 (10x Genomics, California, USA). One-month-old organoids were treated with Cell Recovery Solution (Corning, Cat#354253) for 20 minutes at 4 °C to remove the surrounding Matrigel, and 4-month-old organoids were cut into four pieces and washed with DPBS (Gioco, Cat# 14190144) for three times to remove dead cells from the inner core before cell dissociation with the Neural dissociation kit (P). After the dissociation, the single cell suspensions were filtered with a 100 μm cell strainer (Gibco) followed by 70 μm, and 40 μm Flowmi^®^ cell strainers (Fisher Scientific, Cat#BAH136800070-50EA, Cat# BAH136800040-50EA). An equal number of cells from three separately digested organoids were pooled and loaded onto the 10X Genomics microfluidics chip. The libraries were prepared according to the manufacturer’s protocols and sequenced using the NovaSeq 6000 Paired-End S1 kit (Illumina, California, USA) by the NGS Core Facility of the German Cancer Research Center (DKFZ).

Raw RNA-seq reads were aligned to human genome hg19 (Ensembl v75) with Cell Ranger (v3.1.0) ^59^ with non-default parameter “--expect-cells=10000”. Data from WT, PT, PTCC, and PTN organoids at one- and four-month-old were aligned separately. Raw reads in each condition were analyzed with Seurat (v3.1.5) ^60^. Briefly, cells with the number of features in the quantile range of 5% and 95% in populations, as well as with less than 10% of reads aligned to mitochondrial genes, were used for downstream analysis. We used 75 principal components for dimension reduction, cluster identification, and low-dimension projections. To perform RNA velocity analysis, the splicing information of cells was calculated for each organoid separately with velocyto (v.0.17.17) ^61^. We generated the one- and four-month data by concatenate results across conditions. Looms were converted to h5ad files integrating cell annations and UMAP/t-SNE embedding. RNA velocity was estimated with the stochastic model with generated h5ad files as input to scvelo (v0.2.2) ^62^. The tumor cell state was annotated by mapping the cluster gene signature to the reference cell state signatures ^25^ with Fgsea R package ^63^, the signature with the smallest *P* value was chosen as annotation, in the case when the *P* values were the same, the enrichment score and the gene expression was evaluated to determine the cluster annotation, the cluster remained unmapped if there was no significantly enrichment cell state. The tumor cell meta-module score was calculated with the scallop R package ^9^.

### DNA methylation array and data analysis

The organoids were cut into four pieces and washed with DPBS 3 times to remove the dead cells. The DNA for one-, two- and three-month-old organoids from each genotype was extracted with DNeasy Blood & Tissue Kit (Qiagen, Hilden, Germany, Cat# 69504) following the manufacturer’s protocol. DNA methylation array analyses were then performed with Infinium Methylation EPIC BeadChip Kit (Illumina) according to the manufacturer’s instructions by the microarray unit of the DKFZ Genomics and Proteomics Core Facility.

The DNA methylation EPIC array data were processed with the CHAMP R package ^64^ following the recommended pipeline. A total of 740031 probes were kept for analysis after filtering and normalization. The PCA plot was drawn with the Factoextra R package. Differentially methylated probes (DMP) were identified for PT vs. WT, PTCC vs. WT, and PTN vs. WT at different time points. The differential methylation level of over-methylated and under-methylated probes was calculated by dividing the number of DMP by the total number of probes. The mean delta beta value of all the DMPs localized on specific gene features was used to rank the genes for gene set enrichment analysis (GSEA) ^65^ by the Fgsea R package ^63^. Methylation clustering was performed based on previously identified methylation classification probes ^26^. The MGMT methylation level was calculated by the MGMT-STP27 logistic regression model using the M values of two probes (cg12434587 and cg12981137) ^66^, and the M values were calculated by log transformation of the beta values (M = log_2_(beta/(1-beta)) ^67^. The gene ontology analysis of the stable probes was performed with the clusterProfiler R package ^68^.

### Metabolome and lipidome sample processing and data analysis

#### Sample collection

We performed metabolome and lipidome profiling on one- and four-month-old organoids (five samples in each group) and corresponding culture medium (five samples in each group). Six organoids were transferred to each 6-well-plate well containing 3 mL culture medium and conditioned for two days at 37 °C with 5% CO_2_ on orbital shakers. Blank medium control was prepared by incubating fresh medium under the same condition without organoids. Three organoids were quickly washed with 154 mM ammonium acetate on ice and collected as one sample, and 300 μL medium was collected from each well. All the samples were snap frozen in liquid nitrogen and stored at −80°C before extraction.

#### Organoid extraction (water-soluble metabolites and lipids)

The organoid samples were homogenized with Mixer Mill (Retsch, Haan, Germany) and ceramic beads at maximum frequency for two to four minutes in pre-cooled racks after adding ice-cold methanol/H_2_O (4:1, v/v, 500 μL per 40 mg tissue) with internal standards (4 μM lamivudine, 4 μM D4-glutaric acid, 4 μM D8-phenylalanine, and 16 μl Splash Lipidomix per 40 mg tissue). 500 μL of homogenate was then collected and extracted by applying 60 μL 0.2 M HCl, 200 μL chloroform, 200 μL chloroform, and 200 μL H_2_O consecutively with vortex. The extracts were spun down at 16000 g for ten minutes, and the upper phase (water-soluble metabolites) was evaporated for 30 minutes at 35°C under nitrogen and dried in SpeedVac (Eppendorf, Hamburg, Germany) at 15 °C overnight. The lower phase (lipids) was evaporated to dryness at 45 °C under nitrogen. The interphase was used to determine the protein concentration with the BCA assay. Samples were stored at −80 °C.

#### Culture medium extraction for water-soluble metabolites

The water-soluble metabolites in the culture medium were extracted with RP18 SPE columns (Merck, Darmstadt, Germany, Cat#102014). Briefly, 50 μL medium was mixed with 50 μL H_2_O and 400 μL methanol/acetonitrile (5/3, v/v) containing internal standards (4 μM D4-glutaric acid, 4 μM D8-phenylalanine), vortexed and ultrasound for three minutes. The supernatants were then filtered through the RP18 SPE columns (activated by elution of 1 mL acetonitrile and equilibrated by elution 1 mL methanol/acetonitrile/H_2_O (5/3/2, v/v/v) before usage) after centrifugation (5 minutes, 16000 g, 4 °C). The eluents were collected and mixed with 400 μL of methanol/acetonitrile/H_2_O (5/3/2, v/v/v). The mixtures were vortexed, ultrasound, centrifuged, and filtered as before and the eluent was collected and evaporated in SpeedVac overnight at 15 °C. Samples were stored at −80 °C.

#### Culture medium extraction for lipids

The lipids in the culture medium were extracted with methanol and chloroform. Briefly, 200 μL medium sample was mixed with 800 μL methanol containing internal standards (6 μL Splash Lipidomix). 120 μL 0.2 M HCl, 400 μL chloroform, 400 μL chloroform, and 400 μL H_2_O were added to the mix consecutively and vortexed. The lower phase of the spun-down samples was collected with a 200 μL micro syringe (Hamilton, Reno, USA) and evaporated to dryness at 45 °C under nitrogen. Samples were stored at − 80 °C.

#### LC-MS analysis of water-soluble metabolites

Water-soluble metabolites from organoid and culture medium samples were dissolved in 200 μl 5 mM ammonium acetate (in 75% acetonitrile (v/v)) before loading to LC/MS. LC-MS analysis was performed on an Ultimate 3000 HPLC system (Thermo Fisher Scientific) coupled with a Q Exactive Plus MS (Thermo Fisher Scientific) in both ESI positive and negative mode. The analytical gradients were carried out using an Accucore 150-Amide-HILIC column (2.6 μm, 2.1 mm × 100 mm, Thermo Fisher Scientific) with solvent A (5 mM ammonium acetate in 5% acetonitrile) and solvent B (5 mM ammonium acetate in 95% acetonitrile). 3 μl sample was applied to the Amide-HILIC column at 30°C, and the analytical gradient lasted 20 minutes. During this time, 98% of solvent B was applied for one minute, followed by a linear decrease to 40% within five minutes and maintained for 13 minutes before returning to 98% in one minute and appended with a five-minute equilibration step. The flow rate was maintained at 350 μL/min. The eluents were analyzed with MS in ESI positive/negative mode with ddMS2. The full scan at 70k resolution (69-1000 m/z scan range, 1e6 AGC-Target, 50 ms maximum Injection Time (maxIT)) was followed by a ddMS2 at 17.5k resolution (1e5 AGC target, 50 ms maxIT, 1 loop count, 0.1 s to 10 s apex trigger, 2e3 minimum AGC target, 20 s dynamic exclusion). The HESI source parameters were set as 30 sheath gas flow rate, 10 auxiliary gas flow rate, 0 sweep gas flow rate, spray voltage: 3.6 kV in positive mode, 2.5 kV in negative mode, 320 °C capillary temperature, and the heater temperature of auxiliary gas was 120 °C. The annotation of the metabolites was performed using the EI-Maven software (Elucidata, https://www.elucidata.io/el-maven) with an offset of ± 15ppm.

#### LC-MS/MS analysis of lipids

The lipids from the organoid and culture medium samples were dissolved in 100 μl of isopropylalcohol (iPrOH) before loading. The analytical gradients were carried out using an Accucore C8 column (2.6 μm, 2.1 mm x 50 mm, Thermo Fisher Scientific) with solvent A (acetonitrile/H_2_O/formic acid (10/89.9/0.1, v/v/v)) and solvent B (acetonitrile/iPrOH/H_2_O/formic acid (45/45/9.9/0.1, v/v/v/v)). 3 μl sample was applied to the C8 column at 40°C, and the analytical gradient lasted for 35 minutes. During this time, 20% of solvent B was applied for two minutes, followed by a linear increase to 99.5% within five minutes and maintained for 27 minutes before returning to 20% in one minute and appended with a five-minute equilibration step. The flow rate was maintained at 350 μL/min. The full scan and ddMS2 parameters were the same as the analysis of the water-soluble metabolites, except the scan range were adjusted to 200-1600 m/z. The HESI source parameters were also adapted with a 3-sweep gas flow rate and a 3.2 kV spray voltage in positive and 3.0 kV in negative mode. Peaks corresponding to the calculated lipid masses (± 5 ppm) were integrated using El-Maven software.

#### Metabolome and lipidome data analysis

Two of the four-month-old organoid samples (one in the PT group and one in the PTN group) were removed from downstream analysis due to the low signal intensity of the internal standards. For organoid sample normalization, the intensity of each target was normalized to respective internal standards (positive/negative standards for metabolites, lipid class standards for lipids) and the sample protein concentration. The intensities of medium samples were subtracted by the median of the blank medium before normalizing to internal standards and protein concentrations to visualize the changes driven by organoid metabolism. The metabolomics data was further normalized by variance stabilization normalization (VSN) with the VSN R package ^69^ and significant pathways in the Small Molecule Pathway Database (SMPDB) were identified by the quantitative enrichment analysis with MetaboAnalystR ^70^. The lipidomics data were further normalized by quantile normalization with the Limma R package ^71^, and the enrichment was calculated by comparing the structural similarities with ChemRich ^72^.

### Proteome and phospho-proteome sample processing and data analysis

#### Sample preparation

The proteomics and phospho-proteomics samples (five samples in each group) were prepared according to a previously published protocol with adaptations ^73^. Briefly, cell pellets were resuspended with lysis buffer (100 mM Tris-HCl pH 8.5, 7 M Urea, 1% Triton, 10 U/mL DNase I (, 1 mM magnesium chloride, 1% benzonase, 1 mM sodium orthovanadate, phosphoSTOP phosphatases inhibitors, complete mini EDTA free protease inhibitors) and lysed by sonication. Cell debris was removed by 1.5 hours of 17000 g centrifugation at 4 °C. 1% benzonase was added to the supernatant, followed by incubation at RT for two hours. Protein concentration was determined by the Bradford assay. Proteins were precipitated using chloroform/methanol ^74^, and the pellets were resuspended (8 M Urea, 100 mM NaCl, 50 mM triethylammonium bicarbonate (TEAB), pH 8.5) and reduced in 10 mM dithiothreitol (DTT) for one hour at 27 °C, then alkylated by 30 mM Iodoacetamide for 30 min at RT in the dark and the reaction was quenched by adding additional 10 mM DTT. Samples were subsequently digested by Lys-C at an enzyme: protein ratio of 1:100 for four hours at 30 °C, diluted with 50 mM TEAB to a resulting Urea concentration of 1.6 M, and further digested with Trypsin overnight at 37 °C in an enzyme: protein ratio of 1:50. Digestion was stopped by acidification using 0.02% trifluoroacetic acid (TFA, v/v). Digested peptides were desalted using C18 SepPack Cartridges (Waters) and resuspended in 0.07% TFA (v/v) in 30% acetonitrile (v/v) and fractionated by on-column FE^3+^-Immobilized Metal Ion Affinity Chromatography (IMAC) enrichment on an Ultimate 3000 LC system using the method described previously ^75^. The two resulting fractions per sample, containing either unphosphorylated or phosphorylated peptides, were desalted by StageTips ^76^. Before LC-MS/MS analysis, the dry peptides were resolved in 50 mM citric acid and 0.1% TFA.

#### LC-MS/MS analysis of proteomics

LC-MS/MS analysis was carried out on an Ultimate 3000 UPLC system directly connected to an Orbitrap Exploris 480 mass spectrometer (Thermo Fisher Scientific). Peptides were online desalted on a trapping cartridge (Acclaim PepMap300 C18, 5 μm, 300Å wide pore, Thermo Fisher Scientific) for three minutes using 30 uL/min flow of 0.05% TFA in water. The analytical multistep gradient was carried out using a nanoEase MZ Peptide analytical column (300Å, 1.7 μm, 75 μm x 200 mm, Waters) using solvent A (0.1% formic acid in water) and solvent B (0.1% formic acid in acetonitrile). A total of 150 minutes of LC-MS/MS analysis time was used per sample. The analytical step of the gradient was 134 minutes, during this time, the concentration of B was linearly ramped from 4% to 30% (2% to 28%, for phospho-peptides), followed by a quick ramp to 78%, and after two minutes the concentration of B was lowered to 4% (2% for phospho-peptides) and a 10 min equilibration step appended. Eluting peptides were analyzed with the mass spectrometer using data-dependent acquisition (DDA) mode. A full scan at 120k resolution (380-1400 m/z, 300% AGC target, 45 ms maxIT) was followed by up to 2 seconds of MS/MS scans. Peptide features were isolated with a window of 1.4 m/z (1.2 m/z for phospho-peptides) and fragmented using 26% NCE (28% NCE for phospho-peptides). Fragment spectra were recorded at 15k resolution (100% AGC target, 22 ms maxIT; 200% AGC target, 54 ms maxIT for phospho-peptides). Unassigned and singly charged eluting features were excluded from fragmentation, and dynamic exclusion was set to 35 seconds (10 seconds for phospho-peptides).

#### Target identification and data analysis

Data analysis was carried out by MaxQuant ^77^ (version 1.6.14.0) using an organism-specific database extracted from Uniprot.org under default settings. Identification FDR cutoffs were 0.01 on the peptide level and 0.01 on the protein level. For the phospho enriched fraction, PTM was set to True and Phospho (STY) was added as variable modification. The full proteome samples were given a separate parameter group with the default variable modifications. The match between runs (MBR) option was enabled to transfer peptide identifications across RAW files based on accurate retention time and mass-to-charge ratio. The fractions were set in a condition that MBR was only performed within phospho enriched and full proteome and within each condition. The full proteome quantification was done based on the MaxLFQ algorithm ^78^. A minimum of two quantified peptides per protein was required for protein quantification. LFQ, and phosphosite intensities were filtered for target groups with a non-zero intensity in 70% of the samples of at least one of the conditions and normalized via VSN ^69^. For missing values with no complete absence in one condition, the R package missForest ^79^ was used for imputation. The missing values that were completely absent in one condition were imputed with random values drawn from a downshifted (2.2 standard deviations) and narrowed (0.3 standard deviations) intensity distribution of the individual samples ^80^. The significance for each target was then calculated with Student’s t-test and adjusted with Benjamini-Hochberg method.

The protein abundances of GB subtype signature genes ^42^ were plotted, and the enrichment P value was calculated with the ‘‘ssgsea.GBM. classification” R package ^42^. Enrichment analysis of the full proteome was carried out with the GSEA software (NIH Broad Institute, version 4.2.3) on Hallmark, KEGG, Reactome, and GO biological process gene sets. Potential druggable targets were identified by mapping the significantly differentially expressed proteins (*P* adjusted < 0.05 and foldchange (FC) > 1) to a drug-gene interaction database DGIdb ^43^, and the combinations with an interaction group score more than 5 were kept. The upregulated phospho-sites (FC > 1) were mapped to a kinase/substrate interaction database ^44^ to identify upstream kinases, and the *P* value for each kinase was calculated by Kinase Enrichment Analysis ^45^. The interactions were visualized by Cytoscape ^81^ (version 3.9.1), and the largest subnetwork was shown in the figure.

#### Organoid drug screen

Drug screens on several selected kinase inhibitors (Sellekchem), TMZ (Sigma), one library containing FDA approved drugs that can penetrate through blood-brain barrier (269 drugs from TargetMol, Massachusetts, USA), and one library containing drugs targeting the possible targets identified in omics analysis (58 drugs from MedchemExpress, New Jersey, USA). A list of drug information can be found in Supplementary Table 5.

For drug screening, the organoids generated from Luc2+ iPSCs were cultured with the culture medium containing kinase inhibitors (5 days, 10 μM, applied daily), TMZ (6 days, 100 μM, applied every other day), drug libraries (6 days, 10 μM, applied every other day) or DMSO vehicle on the orbital shakers at 37 °C with 5% CO_2_. BLI was performed before drug administration to one (for daily administration) or two (for every other day administration) days after the last dose. For BLI, the organoids were incubated with 150 μg/mL D-luciferin in a 37 °C incubator supplied with 5% CO_2_ on the orbital shakers for 15 minutes and then imaged with IVIS or Quick View 3000 (Bio Real, Salzburg, Austria). To assess the treatment effects, the BLI signals were normalized to the DMSO control measured on the same day, then compared to before treatment. Drugs with a *P* value less than 0.05 and a signal drop of more than 15% were considered effective. In addition, 100 μM BrdU (Sigma-Aldrich) was applied to the culture medium after the final imaging and cultured for 2 hours, and the samples were collected and stained as described above.

#### Quantification and statistical analysis

All the data analysis was performed with R (version 4.1.2), Graphpad Prism 8 and Microsoft Excel. The 2D areas of the organoids were measured with ImageJ (NIH) and the comparison was carried out with two-way ANOVA. For cell number quantification, positive cells were manually counted using the cell counter function in ImageJ. Group comparisons of Kaplan-Meier survival analysis of xenografted mice were calculated with Log-rank test. Students’ t-tests were applied when comparing variables between two groups (paired t-test for drug treatment and heteroskedastic for the rest). N numbers for each experiment can be found in the corresponding figure legend. All data values were presented as mean ± SEM and the *P* values are represented as follows: **** *P* < 0.0001, *** *P* < 0.001, ** *P* < 0.01, * *P* < 0.05, and *P* > 0.05 is recognized as non-statistically significant (ns).

## Acknowledgments

We thank the DKFZ Next Generation Sequencing Core Facility for providing high throughput sequencing and related services. We acknowledge the DKFZ Microarray Core Facility for providing the Illumina Methylation arrays and associated services. We further thank the team of the DKFZ Proteomics Core Facility for assistance with sample preparation and measurement. Support by the DKFZ Light Microscopy Facility (LMF) and the Discovery Technology Platform at Shanghai Institute for Advanced Immunochemical Studies (SIAIS) of ShanghaiTech University are gratefully acknowledged. We also extend our thanks to Stefan Pusch from the Clinical Cooperation Unit Neuropathology at DKFZ and Ilse Hofmann from the Antibody Core Facility at DKFZ for sharing the NF1 antibody. Finally, we thank the Animal Facility at the National Facility for Protein Science in Shanghai (NFPS), Zhangjiang Lab, China, for providing support in mouse housing and care. C.W. is funded by the China Scholarship Council (CSC) from the Ministry of Education of China. This work was funded by Deutsche Forschungsgemeinschaft grant (SFB1389 to H-K.L.), Deutsche Krebshilfe grant (110227 to H-K.L.), European Research Council (ERC) grant (647055 to H-K.L.), and Deutschen Konsortium für Translationale Krebsforschung (DKTK) grant (AIM2GO to H-K.L.)

## Author contributions

Conceptualization, H-K.L.; Methodology, C.W., M.Sun., C.S., L.S., and Y.Z.; Formal Analysis, C.W., M.Sun, C.S., T.P. and M.Sch.; Investigation, C.W., M.Sun, L.S., Y.Z., Y.H., W.T., N.S., and Y.W.; Resources, L.S., M.Sch., D.M., J-P. M., and A.S.; Visualization, C.W., M.Sun, C.S., and T.P.; Supervision, H-K.L. and A.S.; Project Administration, H.-K.L. and C.W.; Funding Acquisition, H-K.L., and A.S.; Writing – Original Draft, H-K.L., C.W., C.S., M.Sun, M.Sch., D.H.; Writing – Review & Editing, H- K.L., A.S., C.W., Y.H., Y.Z., L.S., D.H.

## Data and materials availability

Any data and information required to reanalyze the data reported in this paper is available from the lead contact upon request. Cell lines generated in this study will be made available with MTA upon request, further information and requests for resources and reagents should be directed to and will be fulfilled by the lead contact Hai-Kun Liu.

## References

1 Moscow, J. A., Fojo, T. & Schilsky, R. L. The evidence framework for precision cancer medicine. Nature Reviews Clinical Oncology 15, 183–192, doi:10.1038/nrclinonc.2017.186 (2018).

2 Hanahan, D. & Weinberg, R. A. The Hallmarks of Cancer. Cell 100, 57–70, doi:10.1016/s0092-8674(00)81683-9 (2000).

3 Letai, A. Functional precision cancer medicine—moving beyond pure genomics. 10.1038/nm0909-1010 23, 1028–1035, doi:10.1038/nm.4389 (2017).

4 Brennan, C. W. et al. The Somatic Genomic Landscape of Glioblastoma. Cell 155, 462–477, doi:10.1016/j.cell.2013.09.034 (2013).

5 McLendon, R. et al. Comprehensive genomic characterization defines human glioblastoma genes and core pathways. Nature 455, 1061–1068, doi:10.1038/nature07385 (2008).

6 Louis, D. N. et al. The 2021 WHO Classification of Tumors of the Central Nervous System: a summary. Neuro-Oncology 23, 1231–1251, doi:10.1093/neuonc/noab106 (2021).

7 Wang, L. B. et al. Proteogenomic and metabolomic characterization of human glioblastoma. Cancer Cell, doi:10.1016/j.ccell.2021.01.006 (2021).

8 Capper, D. et al. DNA methylation-based classification of central nervous system tumours. Nature 555, 469–474, doi:10.1038/nature26000 (2018).

9 Neftel, C. et al. An Integrative Model of Cellular States, Plasticity, and Genetics for Glioblastoma. Cell 178, 835–849.e821, doi:10.1016/j.cell.2019.06.024 (2019).

10 Venteicher, A. S. et al. Decoupling genetics, lineages, and microenvironment in IDH-mutant gliomas by single-cell RNA-seq. Science 355, doi:10.1126/science.aai8478 (2017).

11 Bose, R. & Ma, C. X. Breast Cancer, HER2 Mutations, and Overcoming Drug Resistance. N Engl J Med 385, 1241–1243, doi:10.1056/NEJMcibr2110552 (2021).

12 Chen, J., McKay, R. M. & Parada, L. F. Malignant glioma: lessons from genomics, mouse models, and stem cells. Cell 149, 36–47, doi:10.1016/j.cell.2012.03.009 (2012).

13 Wang, Z. et al. Cell Lineage-Based Stratification for Glioblastoma. Cancer cell 38, 366–379 e368, doi:10.1016/j.ccell.2020.06.003 (2020).

14 Patrizii, M., Bartucci, M., Pine, S. R. & Sabaawy, H. E. Utility of Glioblastoma Patient-Derived Orthotopic Xenografts in Drug Discovery and Personalized Therapy. Front Oncol 8, 23, doi:10.3389/fonc.2018.00023 (2018).

15 Sargent, J. K. et al. Genetically diverse mouse platform to xenograft cancer cells. Dis Model Mech 15, doi:10.1242/dmm.049457 (2022).

16 Kim, J., Koo, B.-K. & Knoblich, J. A. Human organoids: model systems for human biology and medicine. Nature Reviews Molecular Cell Biology 21, 571–584, doi:10.1038/s41580-020-0259-3 (2020).

17 Bian, S. et al. Genetically engineered cerebral organoids model brain tumor formation. 10.1038/nmeth758 15, 631–639, doi:10.1038/s41592-018-0070-7 (2018).

18 Ogawa, J., Pao, G. M., Shokhirev, M. N. & Verma, I. M. Glioblastoma Model Using Human Cerebral Organoids. Cell Rep 23, 1220–1229, doi:10.1016/j.celrep.2018.03.105 (2018).

19 Ying, Q. L. et al. The ground state of embryonic stem cell self-renewal. Nature 453, 519–523, doi:10.1038/nature06968 (2008).

20 Lancaster, M. A. et al. Cerebral organoids model human brain development and microcephaly. Nature 501, 373–379, doi:10.1038/nature12517 (2013).

21 McInnes, L., Healy, J. & Melville, J. UMAP: Uniform Manifold Approximation and Projection for Dimension Reduction. arXiv preprint, arXiv: 1802.03426, doi:10.48550/arxiv.1802.03426 (2018).

22 Paco, A., Aparecida de Bessa Garcia, S., Leitao Castro, J., Costa-Pinto, A. R. & Freitas, R. Roles of the HOX Proteins in Cancer Invasion and Metastasis. Cancers (Basel) 13, doi:10.3390/cancers13010010 (2020).

23 Kim, Y. et al. Perspective of mesenchymal transformation in glioblastoma. Acta Neuropathol Commun 9, 50, doi:10.1186/s40478-021-01151-4 (2021).

24 Varn, F. S. et al. Glioma progression is shaped by genetic evolution and microenvironment interactions. Cell 185, 2184–2199, doi:10.1016/j.cell.2022.04.038 (2022).

25 Johnson, K. C. et al. Single-cell multimodal glioma analyses identify epigenetic regulators of cellular plasticity and environmental stress response. Nature Genetics 53, 1456–1468, doi:10.1038/s41588-021-00926-8 (2021).

26 Sturm, D. et al. Hotspot Mutations in H3F3A and IDH1 Define Distinct Epigenetic and Biological Subgroups of Glioblastoma. Cancer Cell 22, 425–437, doi:10.1016/j.ccr.2012.08.024 (2012).

27 Hegi, M. E. et al. MGMT gene silencing and benefit from temozolomide in glioblastoma. The New England journal of medicine 352, 997–1003, doi:10.1056/NEJMoa043331 (2005).

28 Kim, M. & Costello, J. DNA methylation: an epigenetic mark of cellular memory. Exp Mol Med 49, e322, doi:10.1038/emm.2017.10 (2017).

29 Hanahan, D. Hallmarks of Cancer: New Dimensions. Cancer Discovery 12, 31–46, doi:10.1158/2159-8290.cd-21-1059 (2022).

30 Rusu, P. et al. GPD1 Specifically Marks Dormant Glioma Stem Cells with a Distinct Metabolic Profile. Cell Stem Cell 25, 241–257 e248, doi:10.1016/j.stem.2019.06.004 (2019).

31 Dai, Z., Ramesh, V. & Locasale, J. W. The evolving metabolic landscape of chromatin biology and epigenetics. Nat Rev Genet 21, 737–753, doi:10.1038/s41576-020-0270-8 (2020).

32 Newman, A. C. & Maddocks, O. D. K. Serine and Functional Metabolites in Cancer. Trends Cell Biol 27, 645–657, doi:10.1016/j.tcb.2017.05.001 (2017).

33 Navas, L. E. & Carnero, A. NAD+ metabolism, stemness, the immune response, and cancer. Signal Transduction and Targeted Therapy 6, 2, doi:10.1038/s41392-020-00354-w (2021).

34 Platten, M., Friedrich, M., Wainwright, D. A., Panitz, V. & Opitz, C. A. Tryptophan metabolism in brain tumors - IDO and beyond. Curr Opin Immunol 70, 57–66, doi:10.1016/j.coi.2021.03.005 (2021).

35 Wu, G. et al. Proline and hydroxyproline metabolism: implications for animal and human nutrition. Amino Acids 40, 1053–1063, doi:10.1007/s00726-010-0715-z (2011).

36 Romer, A. M. A., Thorseth, M. L. & Madsen, D. H. Immune Modulatory Properties of Collagen in Cancer. Front Immunol 12, 791453, doi:10.3389/fimmu.2021.791453 (2021).

37 Payne, L. S. & Huang, P. H. The Pathobiology of Collagens in Glioma. Molecular Cancer Research 11, 1129–1140, doi:10.1158/1541-7786.mcr-13-0236 (2013).

38 Sokolowska, E. & Blachnio-Zabielska, A. The Role of Ceramides in Insulin Resistance. Frontiers in Endocrinology 10, 577, doi:10.3389/fendo.2019.00577 (2019).

39 Platten, M., Nollen, E. A. A., Rohrig, U. F., Fallarino, F. & Opitz, C. A. Tryptophan metabolism as a common therapeutic target in cancer, neurodegeneration and beyond. Nat Rev Drug Discov 18, 379–401, doi:10.1038/s41573-019-0016-5 (2019).

40 Liu, Q. et al. The aryl hydrocarbon receptor activates ceramide biosynthesis in mice contributing to hepatic lipogenesis. Toxicology 458, 152831, doi:10.1016/j.tox.2021.152831 (2021).

41 Majumder, S. et al. A genome-wide CRISPR/Cas9 screen reveals that the aryl hydrocarbon receptor stimulates sphingolipid levels. Journal of Biological Chemistry 295, 4341–4349, doi:10.1074/jbc.ac119.011170 (2020).

42 Wang, Q. et al. Tumor Evolution of Glioma-Intrinsic Gene Expression Subtypes Associates with Immunological Changes in the Microenvironment. Cancer Cell 32, 42–56.e46, doi:10.1016/j.ccell.2017.06.003 (2017).

43 Cotto, K. C. et al. DGIdb 3.0: a redesign and expansion of the drug-gene interaction database. Nucleic Acids Research 46, D1068–D1073, doi:10.1093/nar/gkx1143 (2018).

44 Licata, L. et al. SIGNOR 2.0, the SIGnaling Network Open Resource 2.0: 2019 update. Nucleic Acids Research 48, D504–D510, doi:10.1093/nar/gkz949 (2019).

45 Kuleshov, M. V. et al. KEA3: improved kinase enrichment analysis via data integration. Nucleic Acids Research 49, W304–W316, doi:10.1093/nar/gkab359 (2021).

46 Bowman, R. L., Wang, Q., Carro, A., Verhaak, R. G. W. & Squatrito, M. GlioVis data portal for visualization and analysis of brain tumor expression datasets. Neuro-Oncology 19, 139–141, doi:10.1093/neuonc/now247 (2017).

47 Marcus, H. J., Carpenter, K. L., Price, S. J. & Hutchinson, P. J. In vivo assessment of high-grade glioma biochemistry using microdialysis: a study of energy-related molecules, growth factors and cytokines. J Neurooncol 97, 11–23, doi:10.1007/s11060-009-9990-5 (2010).

48 Ferraro, G. B. et al. Fatty Acid Synthesis Is Required for Breast Cancer Brain Metastasis. Nat Cancer 2, 414–428, doi:10.1038/s43018-021-00183-y (2021).

49 Jin, X. et al. A metastasis map of human cancer cell lines. Nature 588, 331–336, doi:10.1038/s41586-020-2969-2 (2020).

50 Vogel, F. C. E. & Schulze, A. Fatty acid synthesis enables brain metastasis. Nat Cancer 2, 374–376, doi:10.1038/s43018-021-00202-y (2021).

51 Orozco, J.M. et al. Dihydroxyacetone phosphate signals glucose availability to mTORC1. Nat Metab 2, 893–901, doi:10.1038/s42255-020-0250-5 (2020).

52 Phillips, H. S. et al. Molecular subclasses of high-grade glioma predict prognosis, delineate a pattern of disease progression, and resemble stages in neurogenesis. Cancer Cell 9, 157–173, doi:10.1016/j.ccr.2006.02.019 (2006).

53 Zhu, Y. et al. Early inactivation of p53 tumor suppressor gene cooperating with NF1 loss induces malignant astrocytoma. Cancer Cell 8, 119–130, doi:10.1016/j.ccr.2005.07.004 (2005).

54 Sakuma, T., Nishikawa, A., Kume, S., Chayama, K. & Yamamoto, T. Multiplex genome engineering in human cells using all-in-one CRISPR/Cas9 vector system. Scientific Reports 4, 5400, doi:10.1038/srep05400 (2015).

55 Brinkman, E. K., Chen, T., Amendola, M. & Bas. Easy quantitative assessment of genome editing by sequence trace decomposition. Nucleic Acids Research 42, e168–e168, doi:10.1093/nar/gku936 (2014).

56 Reuss, D. E. et al. Neurofibromin specific antibody differentiates malignant peripheral nerve sheath tumors (MPNST) from other spindle cell neoplasms. Acta Neuropathol 127, 565–572, doi:10.1007/s00401-014-1246-6 (2014).

57 Lancaster, M. A. et al. Guided self-organization and cortical plate formation in human brain organoids. Nat Biotechnol 35, 659–666, doi:10.1038/nbt.3906 (2017).

58 Lancaster, M. A. & Knoblich, J. A. Generation of cerebral organoids from human pluripotent stem cells. Nat Protoc 9, 2329–2340, doi:10.1038/nprot.2014.158 (2014).

59 Zheng, G. X. Y. et al. Massively parallel digital transcriptional profiling of single cells. Nature Communications 8, 14049, doi:10.1038/ncomms14049 (2017).

60 Stuart, T. et al. Comprehensive Integration of Single-Cell Data. Cell 177, 1888–1902.e1821, doi:10.1016/j.cell.2019.05.031 (2019).

61 La Manno, G. et al. RNA velocity of single cells. Nature 560, 494–498, doi:10.1038/s41586-018-0414-6 (2018).

62 Bergen, V., Lange, M., Peidli, S., Wolf, F. A. & Theis, F. J. Generalizing RNA velocity to transient cell states through dynamical modeling. Nature Biotechnology 38, 1408–1414, doi:10.1038/s41587-020-0591-3 (2020).

63 Korotkevich, G. et al. Fast gene set enrichment analysis (Cold Spring Harbor Laboratory, 2016).

64 Tian, Y. et al. ChAMP: updated methylation analysis pipeline for Illumina BeadChips. Bioinformatics 33, 3982–3984, doi:10.1093/bioinformatics/btx513 (2017).

65 Subramanian, A. et al. Gene set enrichment analysis: A knowledge-based approach for interpreting genome-wide expression profiles. Proceedings of the National Academy of Sciences 102, 15545–15550, doi:10.1073/pnas.0506580102 (2005).

66 Bady, P. et al. MGMT methylation analysis of glioblastoma on the Infinium methylation BeadChip identifies two distinct CpG regions associated with gene silencing and outcome, yielding a prediction model for comparisons across datasets, tumor grades, and CIMP-status. Acta Neuropathologica 124, 547–560, doi:10.1007/s00401-012-1016-2 (2012).

67 Du, P. et al. Comparison of Beta-value and M-value methods for quantifying methylation levels by microarray analysis. BMC Bioinformatics 11, 587, doi:10.1186/1471-2105-11-587 (2010).

68 Yu, G., Wang, L.-G., Han, Y. & He, Q.-Y. clusterProfiler: an R Package for Comparing Biological Themes Among Gene Clusters. OMICS: A Journal of Integrative Biology 16, 284–287, doi:10.1089/omi.2011.0118 (2012).

69 Huber, W., Von Heydebreck, A., Sultmann, H., Poustka, A. & Vingron, M. Variance stabilization applied to microarray data calibration and to the quantification of differential expression. Bioinformatics 18, S96–S104, doi:10.1093/bioinformatics/18.suppl_1.s96 (2002).

70 Chong, J. & Xia, J. MetaboAnalystR: an R package for flexible and reproducible analysis of metabolomics data. Bioinformatics 34, 4313–4314, doi:10.1093/bioinformatics/bty528 (2018).

71 Ritchie, M. E. et al. limma powers differential expression analyses for RNA-sequencing and microarray studies. Nucleic Acids Research 43, e47–e47, doi:10.1093/nar/gkv007 (2015).

72 Barupal, D. K. & Fiehn, O. Chemical Similarity Enrichment Analysis (ChemRICH) as alternative to biochemical pathway mapping for metabolomic datasets. Scientific Reports 7, 14567, doi:10.1038/s41598-017-15231-w (2017).

73 Potel, C. M., Lin, M. H., Heck, A. J. R. & Lemeer, S. Defeating Major Contaminants in Fe(3+)-Immobilized Metal Ion Affinity Chromatography (IMAC) Phosphopeptide Enrichment. Mol Cell Proteomics 17, 1028–1034, doi:10.1074/mcp.TIR117.000518 (2018).

74 Wessel, D. & Flugge, U. I. A method for the quantitative recovery of protein in dilute solution in the presence of detergents and lipids. Anal Biochem 138, 141–143, doi:10.1016/0003-2697(84)90782-6 (1984).

75 Ruprecht, B. et al. 47–60 (Springer New York, 2017).

76 Rappsilber, J., Mann, M. & Ishihama, Y. Protocol for micro-purification, enrichment, pre-fractionation and storage of peptides for proteomics using StageTips. Nature Protocols 2, 1896–1906, doi:10.1038/nprot.2007.261 (2007).

77 Tyanova, S., Temu, T. & Cox, J. The MaxQuant computational platform for mass spectrometry-based shotgun proteomics. Nature Protocols 11, 2301–2319, doi:10.1038/nprot.2016.136 (2016).

78 Cox, J. et al. Accurate proteome-wide label-free quantification by delayed normalization and maximal peptide ratio extraction, termed MaxLFQ. Mol Cell Proteomics 13, 2513–2526, doi:10.1074/mcp.M113.031591 (2014).

79 Stekhoven, D. J. & Buhlmann, P. MissForest--non-parametric missing value imputation for mixed-typem data. Bioinformatics 28, 112–118, doi:10.1093/bioinformatics/btr597 (2012).

80 Tyanova, S. & Cox, J. 133–148 (Springer New York, 2018).

81 Shannon, P. et al. Cytoscape: A Software Environment for Integrated Models of Biomolecular Interaction Networks. Genome Research 13, 2498–2504, doi:10.1101/gr.1239303 (2003).

