## Supplementary information for "A multidimensional atlas of human glioblastoma organoids reveals highly coordinated molecular networks and effective drugs"

### Supplementary figures and tables

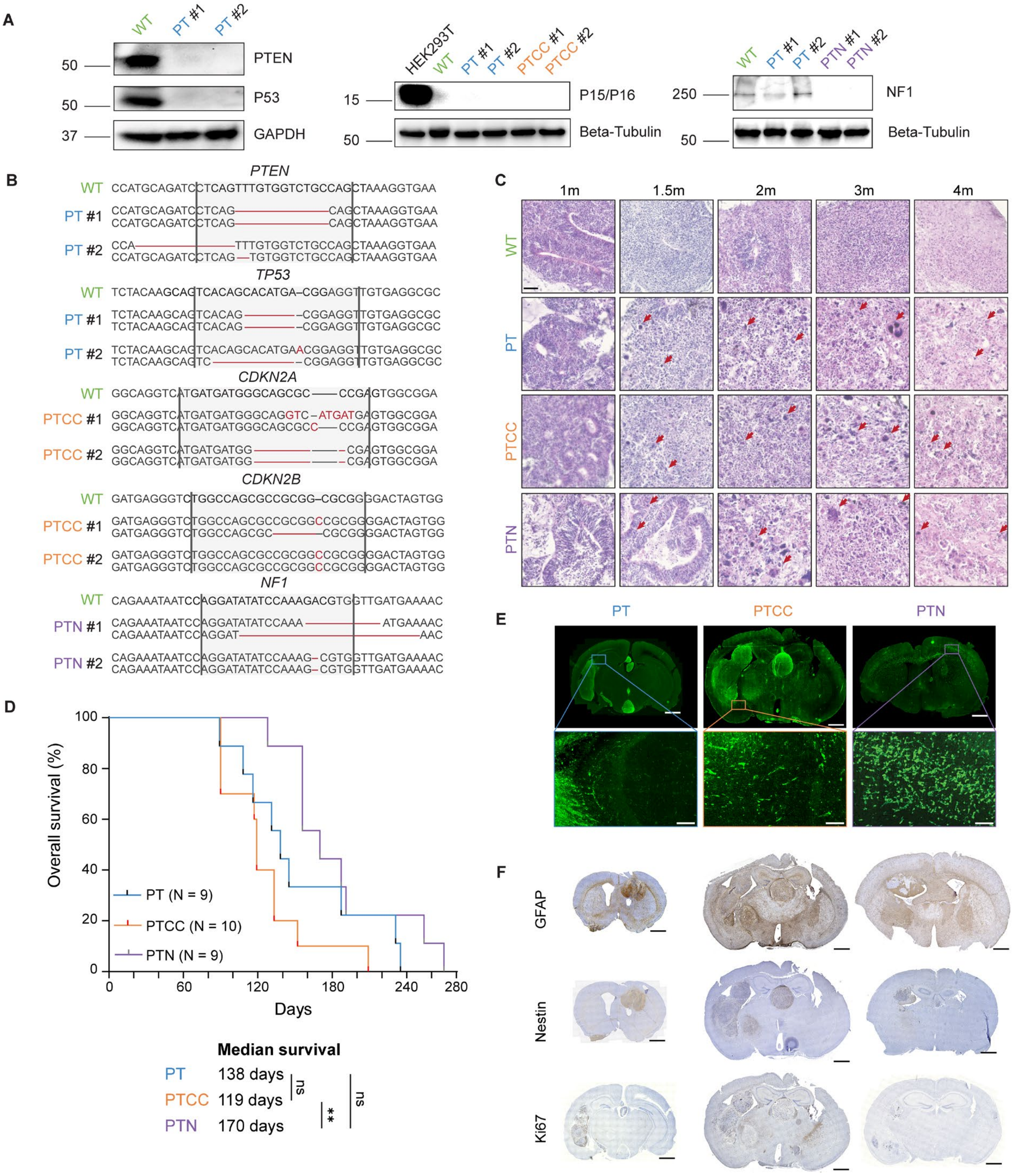


**Figure S1. Related to Figure 1**

(A). Western blotting results of the knockout iPSC clones. HEK293T protein lysate serves as a positive control for P15/P16 detection. The experiments were each performed three times independently.

(B). Sanger sequencing results at the CRISPR/Cas9 gene-editing site of respective iPSC clones. gRNA sequences are highlighted with gray backgrounds, and mutations are highlighted in red.

(C). Representative H&E staining images of the organoids at different ages. Red arrows indicate cells with atypical nuclei. Scale bars, 50 μm.

(D). The Kaplan-Meier survival analysis of LEGO xenografted mice. N numbers of each group are labeled in the figure. P values were calculated with Log-rank test.

(E). Representative GFP staining images showing the infiltrative growth pattern of mouse xenografts. 1000 μm for overview, and 100 μm for insets

(F). Representative IHC staining images with glioma-related markers. Scale bar, 1000 μm.

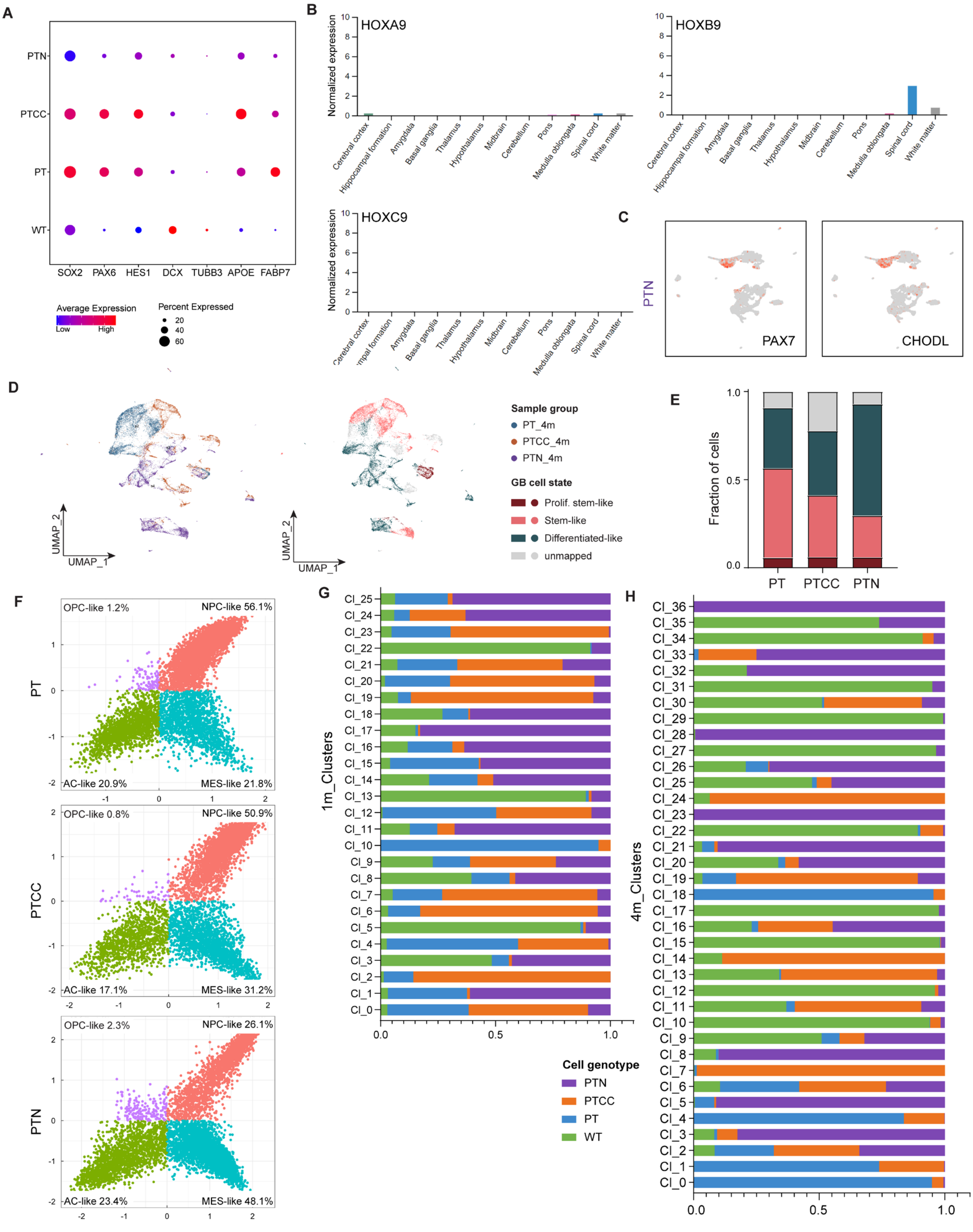

**Figure S2. Related to Figure 2**
(A). Expression portion of lineage markers in different samples.
(B). Normalized RNA expression of HOX genes in different central nervous system regions from the Human Protein Atlas [1].
(C). UMAP gene expression plots of PAX7 and CHODL in four-month-old PTN organoids.
(D). UMAP for four-month-old LEGOs colored by sample (left) and GB cell state (right).
(E). Cell state composition of different LEGOs.
(F). Two-dimensional cellular state annotation with meta-modules from [2].
Genotype composition of each cluster in one- (G) and four-month-old (H) organoids.

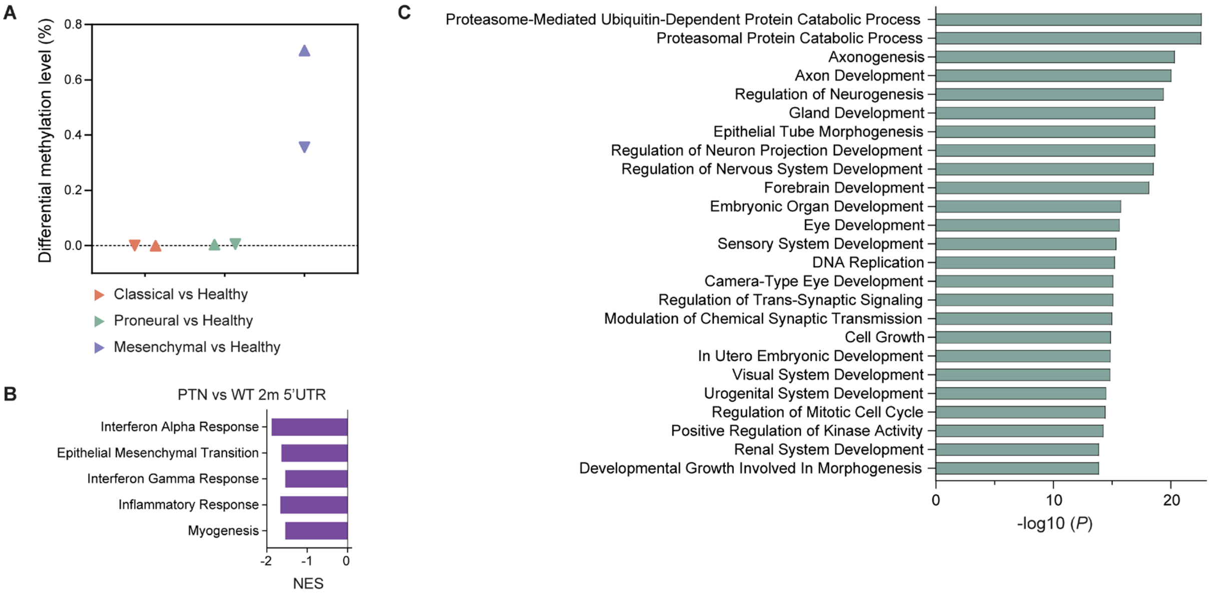


**Figure S3. Related to Figure 3**

(A). Differential methylation level of an external dataset (Sturm et al., 2012) comparing different GBM transcription subtypes to the healthy control.

(B). GSEA hallmark enrichment of the DMPs located on 5'UTR in 2-month-old PTN organoids (adjusted *P* value < 0.05).

(C). The top 25 enriched gene ontology terms of the genes that the stable probes represent.

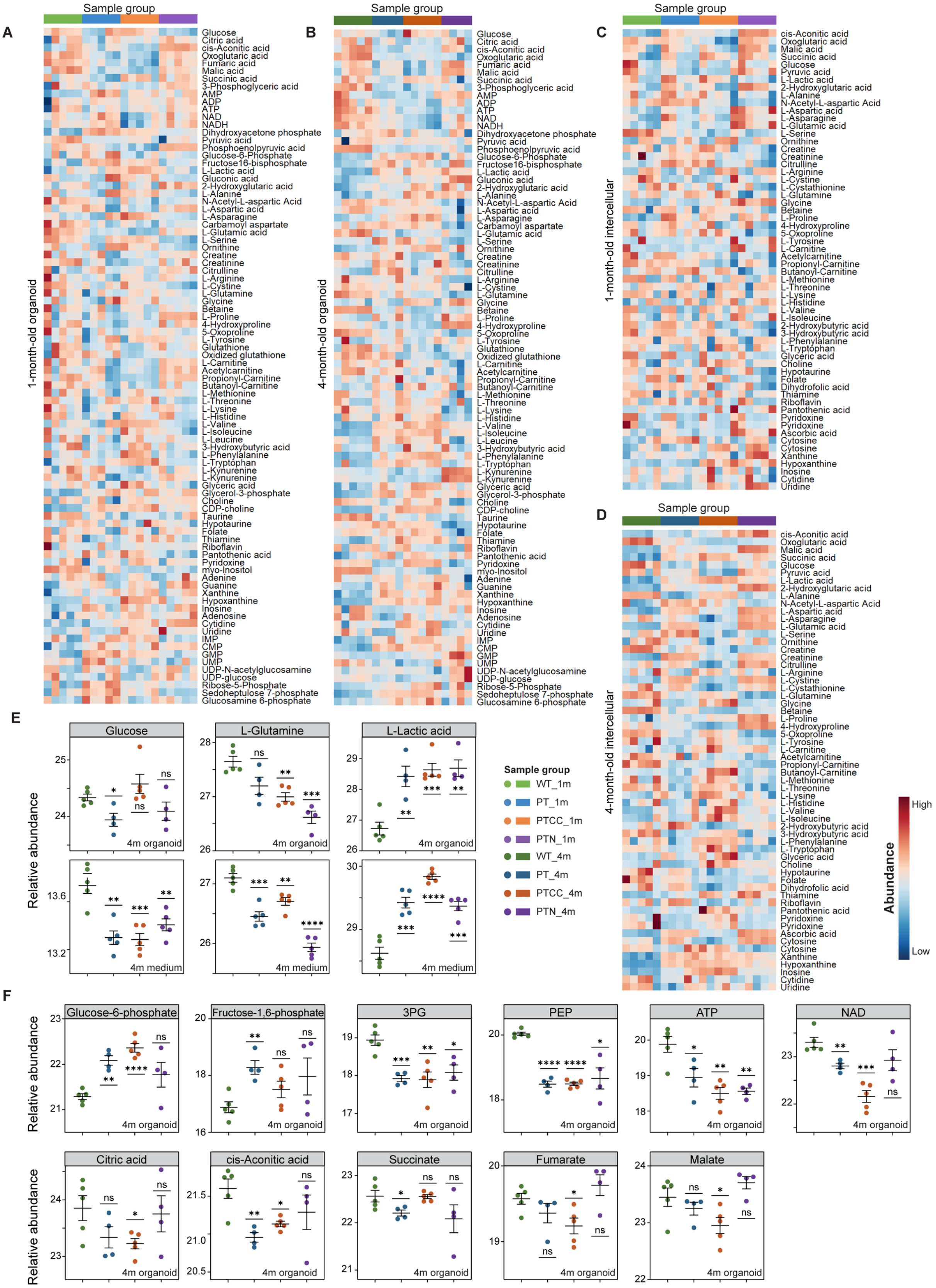

**Figure S4. Related to Figure 4**

(A-D). Heatmap representation of metabolites in one-month-old organoids (A), four-month-old organoids (B), culture medium of one-month-old organoids (C) and culture medium of four-month-old organoids (D).

(E). The relative abundance of energy source (Glucose and Glutamine) and glycolysis product (lactic acid) in four-month-old organoids and culture medium.

(F). The relative abundance of glycolysis and TCA cycle intermediates in four-month-old organoids. 3PG, 3-phosphoglyceric acid; PEP, phosphoenolpyruvic acid.

In E and F, the color of the dots indicates sample group, data are represented as mean ± SEM; N = 4 for four-month-old PT and PTN organoid samples, and N = 5 for the rest of the groups; statistical significances were calculated using Student's t-tests comparing respective mutant groups to WT; **** $P < 0.0001$, *** $P < 0.001$, ** $P < 0.01$, * $P < 0.05$, and ns, non-significant.

A

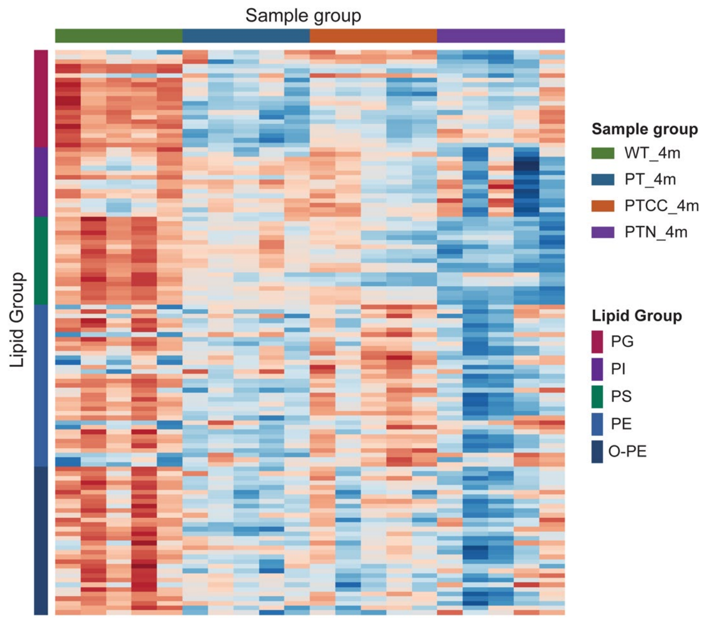


**Figure S5. Related to Figure 5**
(A). Abundance heatmap of structure phospholipid species in four-month-old organoids.

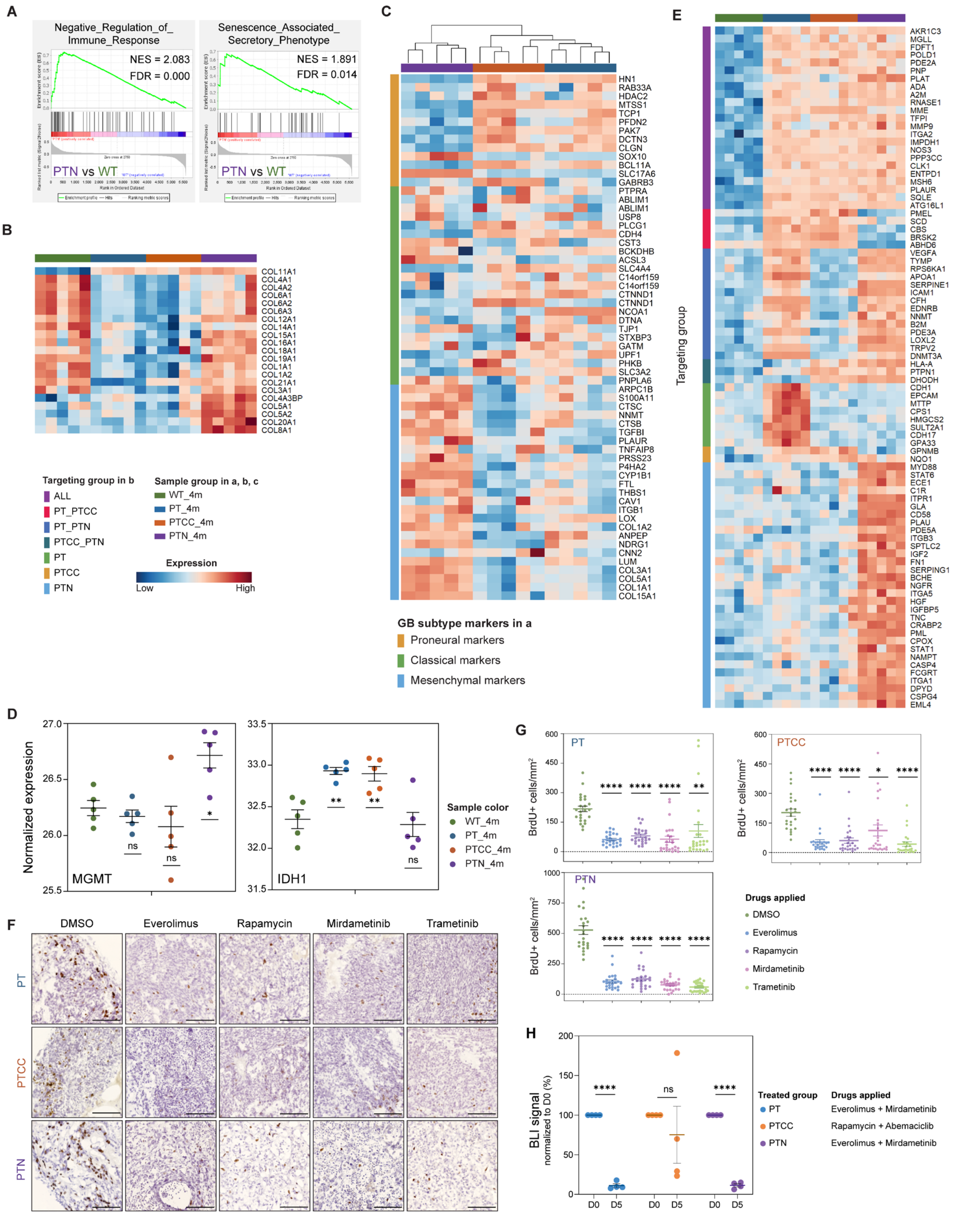


**Figure S6. Related to Figure 6**

(A). GSEA enrichment plots for distinct enriched signatures in PTN.

(B). Heatmap of collagen protein abundance.

(C). Heatmap of protein levels relevant for GB transcriptomic subtype classification [3].

(D). Normalized expression of MGMT and IDH1 protein in different groups of organoids. N = 5 organoids for each group.

(E). Genotype specific drug targets shown in heatmap.

(F). Representative BrdU staining images of LEGOs after drug treatment. Scale bar, 100 μm.

(G). Quantification of BrdU+ cells after drug treatment. N = 24 sections for each group.

(H). Combination treatment in different LEGOs. N = 4 organoids for each group.

In D, G ,and H, the color of the dots indicates the group of organoids; data are represented as mean ± SEM; statistical significances were calculated with Student’s t-test; **** *P* <0.0001, *** *P* < 0.001, ** *P* < 0.01, * *P* < 0.05, ns, non-significant.

**Table S6.** Genotype-based molecular milestones using LEGO model

| | **Tumorigenic *in vivo*** | **scRNA-Seq** | **DNA Methylome** | **Metabolome** | **Lipidome** | **Proteome** |
|---|---|---|---|---|---|---|
| **Shared Features** | Yes | Increased number of stem cells | Dynamic changes in methylation | Phospholipid; Glycolysis; Changes in metabolites regulating methylation (Serine, a-KG, 2-HG) | DG/TG increase; PC increase; Structural lipid decrease | EMT program; Lipid biosynthesis; increase in unfolded protein response |
| **PT** | Growth +; Infiltration +; Angiogenesis + | Astrocyte fate switch + | Methylation change + | | DG/TG increase | AKT1 increase; mTOR increase |
| **PTCC** | Growth ++; Infiltration +; Angiogenesis + | Astrocyte fate switch ++; Activation of WNT pathway | Methylation change ++ MGMT methylated | Abnormal branched-chain amino acid metabolism | DG/TG increase O-PE decrease 1m | AKT1 mTOR CDK1/2/7 increase |
| **PTN** | Growth +; Infiltration +++; Angiogenesis +++ | *HOX* gene activation; EMT increase Mesenchymal cell cluster | Methylation change +++; EMT and inflammation changes | Tryptophan low; Kynurenine high; Proline/hydroxyproline high | Increase of Ceramide; DG/TG increase | Collagen high; Mesenchymal signature; AKT1, mTOR and MAPK increase; IDH1 low; MGMT high |

Abbreviations: α-KG, α-Ketoglutarate; 2-HG, 2-Hydoxyglutaric Acid; DG, Diacylglycerol; TG, Triacylglycerol; PC, Phosphatidylcholine; EMT, Epithelial-Mesenchymal Transition; O-PE, Ether Phosphatidylethanolamine; AKT1, AKT serine/threonine kinase 1; mTOR, Mammalian Target of Rapamycin; CDK, Cyclin Dependent Kinase; MAPK, Mitogen Activated Kinase-like Protein; IDH1, Isocitrate Dehydrogenase 1; MGMT, O6-Methylguanine-DNA Methyltransferase

**Reference**

1 Uhlen, M. *et al.* Proteomics. Tissue-based map of the human proteome. *Science* **347**, 1260419, doi:10.1126/science.1260419 (2015).

2 Neftel, C. *et al.* An Integrative Model of Cellular States, Plasticity, and Genetics for Glioblastoma. *Cell* **178**, 835-849.e821, doi:10.1016/j.cell.2019.06.024 (2019).

3 Wang, Q. *et al.* Tumor Evolution of Glioma-Intrinsic Gene Expression Subtypes Associates with Immunological Changes in the Microenvironment. *Cancer Cell* **32**, 42-56.e46, doi:10.1016/j.ccell.2017.06.003 (2017).
